# Value generalization in human avoidance learning

**DOI:** 10.1101/223149

**Authors:** Agnes Norbury, Trevor W. Robbins, Ben Seymour

## Abstract

Generalization during aversive decision-making allows us to avoid a broad range of potential threats following experience with a limited set of exemplars. However, over-generalization, resulting in excessive and inappropriate avoidance, has been implicated in a variety of psychological disorders. Here, we use reinforcement learning modelling to dissect out different contributions to the generalization of instrumental avoidance in two groups of human volunteers (*N*=26, *N*=482). We found that generalization of avoidance could be parsed into perceptual and value-based processes, and further, that value-based generalization could be subdivided into that relating to aversive and neutral feedback - with corresponding circuits including primary sensory cortex, anterior insula, and ventromedial prefrontal cortex, respectively. Further, generalization from aversive, but not neutral, feedback was associated with self-reported anxiety and intrusive thoughts. These results reveal a set of distinct mechanisms that mediate generalization in avoidance learning, and show how specific individual differences within them can yield anxiety.

## Introduction

During aversive decision-making, generalization allows application of direct experience with a limited subset of dangerous real-world stimuli to a much larger set of potentially related stimuli. For example, if eating a particular foraged fruit has led to food poisoning in the past, it may be adaptive to avoid similar-appearing fruit in the future. As an evolutionarily well-conserved process, generalization enables safe and efficient navigation of a complex and multidimensional world (Sutton and Barto, 1998; Ghirlanda and Enquist, 2003). However, over-generalization, resulting in inappropriate avoidance of safe stimuli, actions or contexts, and has been suggested as a possible pathological mechanism in a range of psychological disorders including anxiety, chronic pain, and depression (Duits et al., 2015; Dymond et al., 2015; Vlaeyen and Linton, 2012; Harvie et al., 2017; Pearson et al., 2015).

Previous work on aversive generalization has focused on predicting punishments in passive (Pavlovian) designs. Such studies have revealed evidence of heightened subjective, physiological and neural responses to stimuli that bear perceptual similarity to learned exemplars (Dymond et al., 2015). However, the extent to which these observations extend to a decision-making context - i.e. whether or not to make an avoidance response in the face of certain stimuli, allowing us to exert *control* over experience of aversive outcomes - is unclear. Although Pavlovian processes can influence avoidance learning, the latter involves acquisition of a fundamentally distinct set of values relating to actions themselves. This is a clinically important distinction, as theories of many psychological disorders relate specifically to excessive avoidant behaviour *over and above* subjective fear (Krypotos et al., 2015) - for example, by reducing opportunities for extinction of inappropriate fear or allowing unnecessary avoidance to transfer to habit-based control (Arnaudova et al., 2017; LeDoux et al., 2017; Gillan et al., 2014).

There are a number of potential mechanisms by which avoidance generalization could be implemented by the brain. As emphasised in some accounts, perceptual uncertainty in stimulus identity alone can effectively yield generalization. Although there is debate about how well discriminative ability is controlled for in many generalization experiments (Struyf et al., 2015), there is good evidence that experience with aversive outcomes alters the representation of predictive stimuli in primary sensory cortices (Weinberger, 2007; Sasaki et al., 2010; Wigestrand et al., 2017), and that this may result in changes to absolute stimulus discriminability (Resnik et al., 2011; Laufer and Paz, 2012; Aizenberg and Geffen, 2013). On the other hand, generalization may also occur at the level of *value* representations, by the transfer of acquired value to similar, but discriminable cues during learning. In the Pavlovian case, several well-established behavioural phenomena implicate value-related processes at play in generalization across species (Hanson, 1959; Schechtman et al., 2010). That both perceptual and value processes might operate in parallel may explain why recent neuroimaging studies have highlighted different brain areas (e.g. limbic cortex vs primary sensory regions) as being key to Pavlovian aversive generalization in humans (Onat and Büchel, 2015; Laufer et al., 2016).

A further important factor in the control of avoidance learning is reinforcement by neutral (or ‘safety’) states, that signal omission of punishment. It is likely that generalization over these states can also influence behaviour: for example in the Pavlovian case, evidence for this is seen in ‘peak-shift’ effects, whereby the presence of a local safety cue appears to inhibit response to nearby aversive cues (Hanson, 1959). It is therefore possible that under-generalization of safety cues, as opposed to over-generalization of aversive cues, might be a contributing factor to susceptibility to disorders such as generalized anxiety in humans (Grupe and Nitschke, 2013).

Here, we address three key questions: first, is there good evidence for generalization in avoidance learning in humans?; second, can we distinguish behavioural and neural components relating to perceptual, aversive value, and safety value?; and third, which if any component predicts relevant self-reported psychological symptoms? We used a custom-designed perceptual task in conjunction with reinforcement learning modelling to study two groups: a laboratory-based sample (N=26) who performed a pain avoidance task with concurrent neuroimaging (fMRI), and a larger cohort of individuals (N=482), who performed a monetary loss avoidance task online alongside a battery of questionnaires designed to probe relevant psychological symptom dimensions (Gillan and Daw, 2016).

## Results

The overall study design is summarised in **Figure 1a**. In both groups of participants, generalization of instrumental responding was tested using a costly avoidance paradigm **(Figure 1c)**. Briefly, participants were instructed that they would see a series of flower-like shapes on their screen, some of which were ‘safe’, and some of which were ‘dangerous’. If they saw a dangerous shape and made no response, there was a high chance that they would receive a painful electric shock (fMRI sample), or lose 10 cents from their cash stake (online sample using Amazon Mechanical Turk, AMT). If they saw a safe shape, they would never receive a shock (or lose money) on that trial. In order to escape the possibility of a painful shock (or monetary loss) when they thought a dangerous shape had been presented, participants were told they could press the ‘escape’ button on their keypad. Participants were instructed that the aversive outcome would never occur on a trial when they had pressed the ‘escape’ button – but – that, importantly, pressing the button was associated with a small cost. Specifically, each time they pressed the escape button, it would be registered on a counter at the bottom of their screen. At the end of each block of the task, they would receive additional painful shocks (or lose additional cash) depending on how many times they had pressed the button during that block (one extra shock or 10 cent loss per every 5 button presses). The optimal strategy (in order to minimise the amount of pain received or money lost) would therefore be to press the button if they thought they saw a dangerous shape, but *not* press if they thought a safe shape was on the screen.

**Figure 1:**
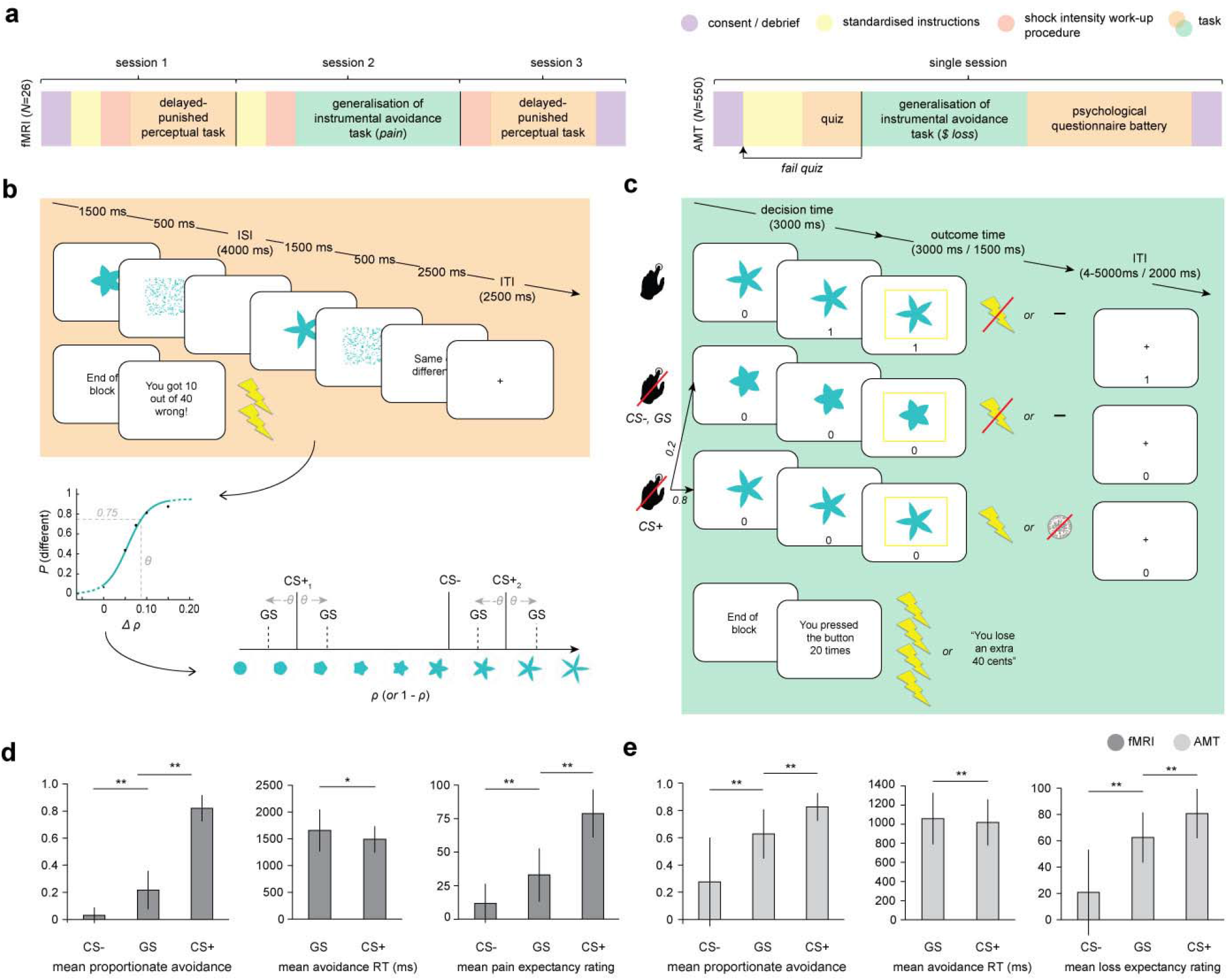
Study design and overall behaviour summary. **a**, Study design and protocol for the two participant groups; fMRI, laboratory and functional imaging sample; AMT, Amazon Mechanical Turk (web-based) sample. **b**, Delayed-punished perceptual task, used to determine 75% reliably perceptually distinguishable generalization stimuli (GSs) on in individual basis for the generalization of instrumental avoidance task (**c**) in the fMRI sample (in the AMT sample, GSs were generated based on mean perceptual acuity determined in pilot testing). **d**, Summary of behaviour on the generalization task in fMRI and **e**, AMT samples. ISI, inter-stimulus interval; ITI, inter-trial interval; CS+, conditioned stimulus with pain or loss outcome, CS-, conditioned stimulus with neutral outcome (no pain or loss). Error bars represent SD. \**p*=0.006, \*\**p*<0.001, paired sample t-tests.

Crucially, on a small proportion of trials, the presented shapes were generalization stimuli (GSs). GSs were individually generated using precise estimates of perceptual ability (as measured on the first study session for the fMRI group) to be 75% reliably perceptually distinguishable from the task stimuli associated with aversive outcomes (CS+s). (Due to time constraints and lack of control over testing environment, GS were generated based on average perceptual acuity from a pilot study in the online group.) The perceptual task **(Figure 1b)** was custom designed based on the recommendations of a recent review (Struyf et al., 2015). Specifically, in order to provide a fair test of perceptual performance during the generalization task, stimuli were not instantly comparable (in order to ensure that GSs would be reliably discriminable in an absolute sense, when presented in isolation; Slivinske and Hall, 1960), and testing occurred in the same emotional context (i.e. under threat of painful shock).

Importantly, the task stimulus array (in terms of arrangement of CS+ and CS- stimuli in perceptual space) was specifically chosen to probe asymmetries in generalization behaviour that result from value-based mechanisms – see **Figure 1b**. One such potential asymmetry is a characteristic shift in peak responding from the CS+ to surrounding GSs, away from the direction of the CS- in perceptual space (known as ‘peak shift’), that has been proposed to result from the interaction of excitatory and inhibitory generalization gradients around CS+ and CS- stimuli following Pavlovian conditioning (Hanson, 1959). Crucially, the asymmetric array used here allowed us to compare responses to CS+ GSs both near and far in perceptual space from the CS- – enabling detection of gradient interaction effects such as peak shift in instrumental avoidance, and allowing the separation of oppositely signed generalization gradients around CS+ and CS- stimuli.

## Evidence for generalization in avoidance behaviour

For both groups of participants, the frequency of avoidance in response to generalization stimuli was intermediate to that evoked by CS- and CS+ stimuli (all *p*<0.0001, paired-sample *t* tests; fMRI: GS vs CS- *t_25_*=7.57, mean difference=0.18 [95%CI 0.14-0.24], GS vs CS+ *t*_25_=-17.6, mean difference=-0.60 [95%CI -0.67-0.54]; AMT: GS vs CS- *t*_481_=27.0, mean difference=0.35 [95%CI 0.33-0.38], GS vs CS+ *Usi=-* 26.6, mean difference=-0.20 [95%CI -0.19—0.21]; **Figure1d,e**). Despite never having been associated with the aversive outcome, participants also rated GSs significantly higher than CS- (but lower than CS+) stimuli on post-task pain/loss expectancy scales (all *p*<0.0001, paired-sample *t* tests; fMRI: GS vs CS- *t*_25_=5.69, mean difference=24.1 [95%CI 15-33], GS vs CS+ *t*_25_=-8.14, mean difference=-52 [95%CI -39−-66]; AMT: GS vs CS- *t*_481_=29.4, mean difference=41.7 [95%CI 40.0-44.6], GS vs CS+ *t*_481_=-16.5, mean different =-18 [95%CI -16.0–-20.3], on visual analogue scales ranging 0-100; **Figure1d,e**).

There was also a significant positive relationship between relative GS avoidance and relative GS pain/loss expectancy rating post-task in both groups (fMRI, Spearman’s *ρ*=0.655, *p*=0.00027; AMT, Spearman’s *ρ*=0.432, *p*=2.2e-16; both measures within-participant z-transformed, for relationships between raw scores see **Figure S1**). This suggests that a higher frequency of avoidance responding (plus associated lack of extinction) translated into higher conscious negative expectancy beliefs for generalization stimuli. There was no relationship between proportionate avoidance on GS trials and perceptual acuity at session 1 (individual θ values) or absolute intensity of the painful electrical stimulation (current amplitude) in the fMRI sample (all p> 0.2).

This raises the question as to whether the observed avoidance on the GS trials was over and above that which would be expected from perceptual uncertainty alone. Notably, mean proportionate avoidance on GS trials in the fMRI group was around 0.2 (or~0.25 when scaled relative to individual CS+ avoidance) – which, given that GSs were generated to be 75% reliably distinguishable from CS+s, is what might have been predicted from a purely perceptual account of task performance. Mean reaction times for making avoidance responses were also significantly slower for GS compared to CS+ stimuli in both groups, suggesting greater uncertainty on these trials (*p*=0.006, *p*=2.07e-11, paired sample *t* tests; fMRI: *t*_25_=3.00, mean difference=167ms [95%CI 51.2-282], AMT: *t*_481_=6.87, mean difference=38.8ms [95%CI 27.7-49.9]; **Figure1d,e**). To resolve this issue, we tested for the presence of additional value-based generalization processes in both datasets using a principled model comparison approach.

Simply, we fitted a series of reinforcement learning models to avoidance data from both samples (modified SARSA algorithms, with trial-by-trial varying learning rates determined by the Pearce-Hall associability rule; Sutton and Barto, 1998; Pelley, 2004-see Methods). Firstly, we fit a model with perceptual ‘generalization’ only (modelled as 25% chance of perceptual confusion between GSs and the adjacent CS+). Secondly, we fit a model with perceptual generalization plus an additional value-based generalization process. As there is some evidence that generalization functions are approximately Gaussian in shape, at least along a single perceptual dimension (Ghirlanda and Enquist, 2003), this was implemented as a Gaussian smoothing of stimulus value across perceptual space, with a single free parameter (σ) governing the width of this function. Thirdly, we fit a model with perceptual generalization plus two additional free parameters governing width of additional value-based generalization processes – one for aversive (shock/loss) and one for neutral (no shock/no loss) feedback (σ_A_ and σ_N_, respectively). This model was informed by previous empirical observations that generalization functions vary in gradient or width for aversive, neutral, and rewarding feedback (Schechtman et al., 2010; Resnik and Paz, 2015; Laufer et al., 2016).

The above models were fit to avoidance data from both groups using a variational Bayes approach to model inversion, under a mixed-effects framework (whereby within-subject priors are iteratively refined and matched to the inferred parent population distribution; see Methods). Random-effects Bayesian model comparison indicated that in both samples the model with two additional value-generalization mechanisms (separately governing width of generalization from aversive and neutral feedback) best accounted for the avoidance data, as indexed by exceedance probability (probability that the model in question was the most frequently utilised in the population; fMRI, EP=0.823, AMT, EP=~1; **Figure 2a**).

**Figure 2:**
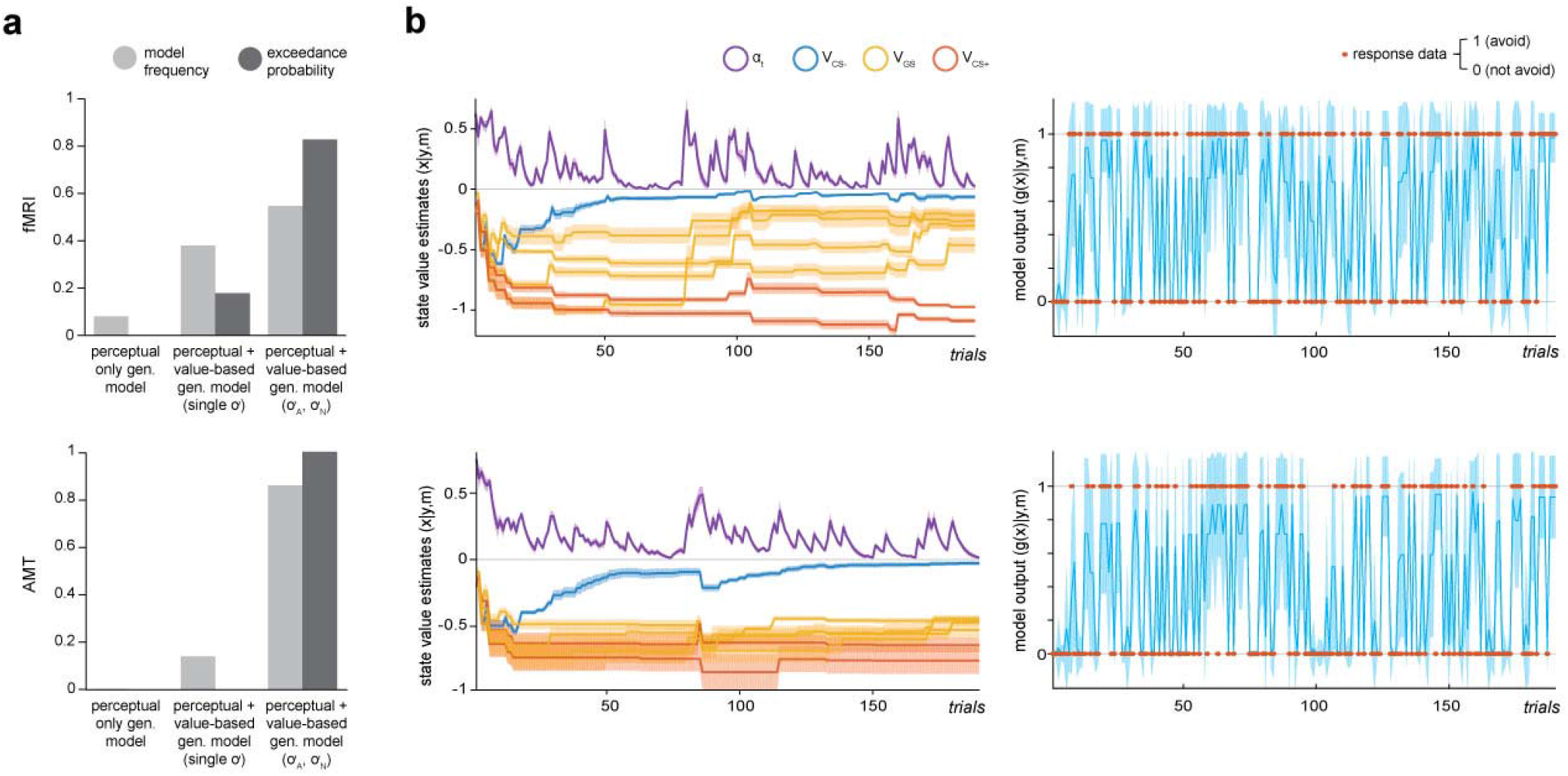
Computational modelling of instrumental avoidance behaviour. **a**, Results of random-effects Bayesian model comparison for the laboratory (fMRI) and online (AMT) samples. For both groups, the best model was one that implemented both perceptual and additional value-based generalization between stimuli, with separate parameters governing width of generalization from aversive (σ_A_) and neutral (σ_N_) feedback. Model frequency, proportion of participants for whom a model was the best model; exceedance probability, probability that the model in question is the most frequently utilized in the population, **b**, llustration of posterior state value estimates (x; here this is the value of not avoiding for each CS, V_cs_, plus the trial-varying learning rate, α_t_) and model output (g(x)) for the winning model (m) for a lower generalizing participant (top row) and higher generalizing participant (bottom row) from the fMRI group. Orange dots on the right hand side panels illustrate actual response data (y) on each trial. Shading represents variance of the posterior density.

For both fMRI and AMT data, this model provided a good account of avoidance choices. Mean predictive accuracy (*r*^2^, for binary choice data this is equivalent to the percentage of correct classifications) was 0.868 (± 0.07) for fMRI and 0.849 (± 0.11) for AMT groups, and the Bayesian ‘p value’ (posterior probability of the null hypothesis of random choice) was ≤6.8e-7 for all fMRI participants, and ≤0.026 for 477/482 AMT participants. In both groups, values of the parameter describing the width of aversive feedback (σ_A_) were unrelated to values of other model parameters governing learning rate, choice bias, and choice stochasticity (see Methods; all p>0.09), suggesting sufficient parameter identifiability. In both samples, σ_A_ values were significantly larger than values of the parameter governing width of generalization from neutral (safe) feedback, σ_N_, indicating wider generalization for aversive compared to neutral outcomes (*p*<3.0e-8, *p*<2.2e-16, related-samples Wilcoxon signed rank tests; fMRI: mean σ_A_=0.752 ± 0.29, mean σ_N_=0.028 ± 0.03; AMT: mean σ_A_=0.695 ± 0.23, mean σ_N_=0.057 ± 0.05). Interestingly, σ_A_ values were not significantly related to σ_N_ values (fMRI group, Spearman’s ρ=-0.169, *p*>0.4; AMT group, ρ=0.06, *p*>0.17), suggesting these may be at least partially independent processes.

Importantly, only a model including additional value-based generalization mechanisms can generate asymmetries in avoidance behaviour across pairs of generalization stimuli (peak shift), as apparent in **Figure S2**. Further, example traces for two representative participants from the fMRI group **(Figure 2b)** illustrate that stimulus values tend to asymptote – i.e. that under this model generalization of value across stimuli is assumed to be relatively constant over time. This assumption is consistent with our behavioural data, in that a time-on-task analysis showed that after initial period of exploratory learning (blocks 1-2), generalization in terms of GS avoidance remains fairly stable **(Figure S2,** see Supplementary Results for a full time-on-task analysis of avoidance data in both groups).

## Evidence for effects of conditioning on perceptual acuity

In the fMRI group, perceptual acuity for task stimuli was tested both before and after carrying our the generalization of instrumental avoidance paradigm, in order to test for possible effects of aversive conditioning on discriminability of the generalization stimuli (the three test sessions were carried out on three consecutive days for all participants, so any detected changes would likely reflect postconsolidation changes in perceptual performance).

There was no strong evidence for change in perceptual acuity in terms of θ value (difference in shape ‘spikiness’ parameter rho for 75% reliable perceptual discrimination) pre- vs post-conditioning (mean θ 0.071 ± 0.015 on session 1, 0.065 ± 0.019 on session 3; non-significant trend towards greater acuity on session 3, p=0.061, related-samples Wilcoxon signed rank test; **Figure S3**). Bayesian model comparison indicated that a model where generalization stimulus discriminability was held constant at 75% better accounted for avoidance data than one where discriminability was held constant at the estimated post-test (session 3) level, or a model where GS discriminability was assumed to be linear between session 1 and session 3 values (exceedance probability for the 75% constant model=^~^1; **Figure S3**). Therefore GS discriminability was held constant across trials at 75% in all models.

## Brain regions encoding model quantities specific to value-based generalization

As our behavioural data provided evidence for the presence of generalization in instrumental avoidance in both groups, we next employed a univariate analysis approach to our functional imaging data in order to investigate whether model quantities specific to *value*-related generalization processes were encoded in regional blood oxygen level-dependent (BOLD) signals.

In addition to work highlighting the role of the insula, amygdala, and primary sensory cortex in aversive generalization following Pavlovian conditioning (Resnik and Paz, 2015; Ghosh and Chattarji, 2015; Onat and Büchel, 2015; Laufer et al., 2016), previous functional imaging studies have identified the striatum and prefrontal cortex as encoding generalization gradients in healthy human volunteers (Dunsmoor et al., 2011; Greenberg et al., 2013; Lissek et al., 2014). However, the contribution of perceptual uncertainty (i.e. absolute discriminability of ‘generalization stimuli’ compared with other conditioned stimuli) is not always adequately addressed in the study of such gradients. Here, we used a strict parametric approach to identify additional variance in regional BOLD that can be attributed to our winning value-based generalization model, *over and above* that which can be explained by a purely perceptual account. This was achieved by using serially orthogonalised regressors derived from each model to predict trial-by-trial variation in BOLD signal in our regions of interest (see **Figure 3a** and Methods).

**Figure 3:**
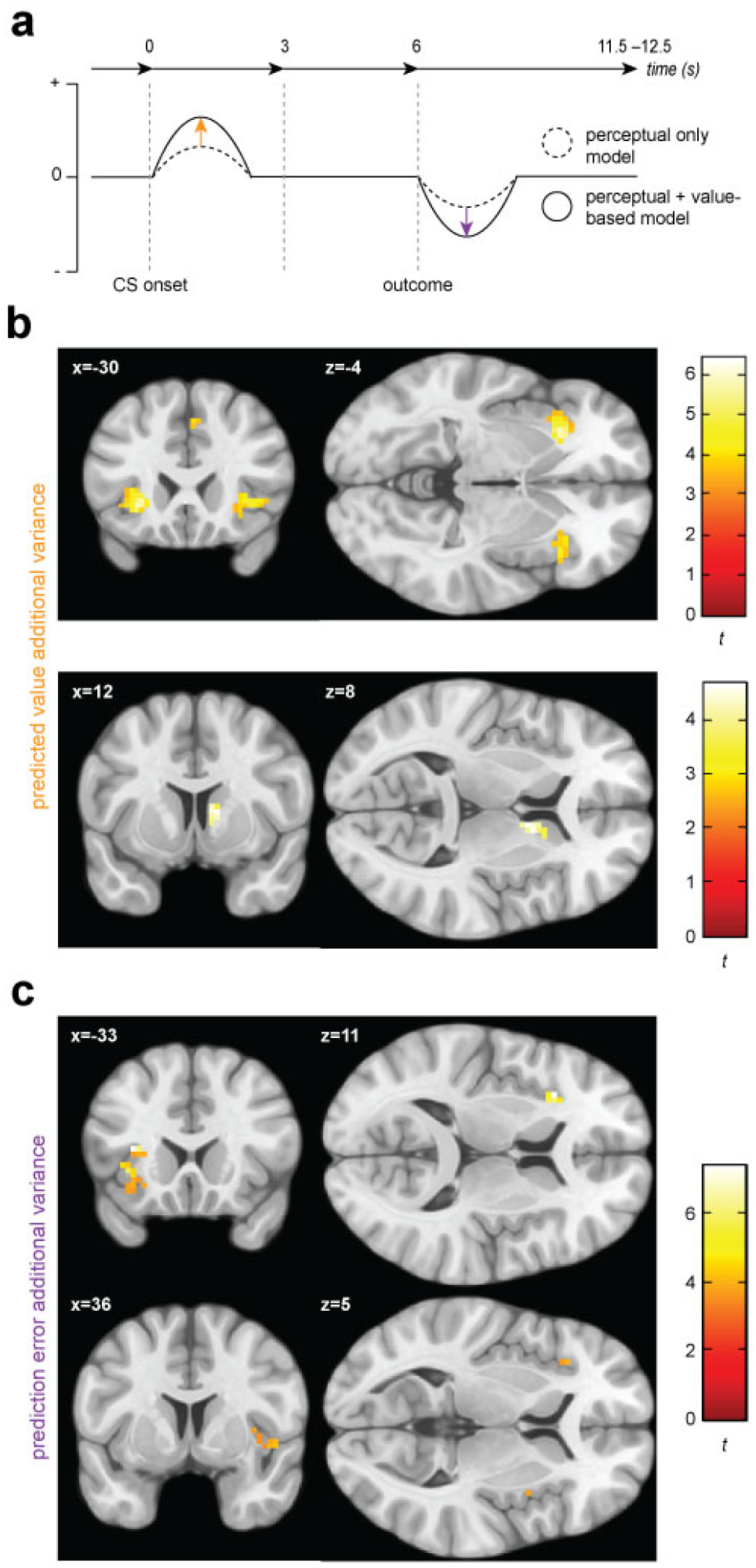
Univariate statistical maps highlight brain regions where changes in BOLD signal is significantly related to trial-by-trial variance in internal model quantities from the value-based generalization model, over and above that which can be explained by a purely perceptual account. **a**, Schematic of a single trial for the fMRI group, showing the difference in estimated probability of receiving a shock (if no avoidance response is made) and prediction error for a generalization stimulus, as derived from the perceptual only vs the perceptual + additional value-based generalization models. **b,** Significant encoding of additional value-based generalization in the expected value of each stimulus (likelihood of receiving a painful shock if no avoidance response is made), at the time of stimulus onset in the anterior insula, and right caudate. **c,** Significant encoding of additional value-based generalization as expressed in prediction error at time of outcome receipt in the left anterior insula, and right insula, more posteriorly.

We found evidence for the encoding of additional variance in trial-by-trial expected stimulus values derived from the value-based generalization model in both the anterior insular cortex and the dorsal striatum **(Figure 3b)**. The BOLD signal was greater when the expected value of a particular stimulus was lower (or the predicted probability of receiving a painful shock if an avoidance response was not made was higher) in the anterior insula, bilaterally (left anterior insula: *p*_WB_=0.0037, *k*=88, peak voxel [-30,23,-4], *Z*=4.57; right anterior insula: *p*_WB_=0.088, *p*_SVC_=0.015, *k*=40, peak voxel [33,20,5], *Z*=3.92), and the right caudate (*p*_SVC_=0.015, *k*=26, peak voxel [12,8,8], *Z*=3.93).

There was also a trend towards significant encoding of value generalization-derived expected value signals in the mid cingulate cortex (Supplementary Motor Area: *p*_WB_=0.070, *k*=A3, peak voxel [9,14,44], *Z*=4.88), which may reflect motor preparation for avoidance responses (see Methods) – but no evidence for univariate encoding of this signal in primary visual cortex (V1) or the amygdala. We also found no evidence for *negative* encoding of aversive value (i.e. greater BOLD signal with lower predicted probability of shock i.e. ‘safety signalling’) in the ventromedial prefrontal cortex (vmPFC).

In addition to expected value signals, we examined potential encoding of prediction errors, which are the main learning signals in reinforcement learning (PEs; defined as the difference between actual and predicted outcome on any given trial – see Methods). We focused our analysis on negatively signed PEs (generated on trials where no shock was received, but the predicted *P*(shock) was >0), as this both constrains analysis to trials where an avoidance response was not made (on avoidance trials PE=0, by definition), minimising potential contamination by motor preparation responses, and gives greater weighting to generalization trials where, due to perceptual uncertainty alone, predicted P(shock) will be >0, but no aversive outcome is ever delivered. (Positively signed PEs are highly collinear with shock administration and therefore are hard to detect under our design.)

We found evidence of significant encoding of additional variance in PE signals from the value-based generalization model in the insula and inferior parietal lobule **(Figure 3c)**. Specifically, BOLD signal was greater when trial PE was lower in the left anterior insula (*p*_WB_=0.026, *k*=55, peak voxel [-33,20,11], Z=5.34), the right insula more posteriorly (*p*_WB_=0.030, *k*=43, peak voxel [48,8,-4], *Z*=4.07), and the right inferior parietal lobule (*p*_WB_=0.026, *k*=55, peak voxel [54,-40,26], *Z*=3.80). There was a non-significant trend in the same direction in the right putamen (*p*_SVC_=0.074, *k*= 10, peak voxel [33,2,-1], *Z*=3.47), but no evidence of encoding of value generalization-derived PE signals in V1, the amygdala, or vmPFC.

## Changes in neural representation of generalization stimuli over the course of the task and relationship to individual differences in avoidance behaviour

Previous evidence from animal research has shown that over the course of conditioning, the representation of the conditioned stimulus (CS+) in terms of response pattern across many individual units may come to resemble that of the primary aversive reinforcer (e.g. Grewe et al., 2017). To complement our univariate results, we therefore examined how different task stimuli were represented in multivariate space using representational similarity analysis (Kriegeskorte et al., 2008). This approach enables the consideration of the full representational geometry across specific brain regions – *how* information is encoded, as well as whether or not it is – and depends on the calculation of distance metrics to quantify how (dis) similarly different kinds of stimuli are represented in multivariate space (in fMRI, across all voxels in a particular brain volume).

Following the approach of a recent electrophysiological study of aversive conditioning in rodents (Grewe et al., 2017), we examined how representational difference changed in our regions of interest earlier (blocks 1-2) vs later (blocks 3-5) in the task – and, crucially, how this change related to individual differences in overall behavioural expressions of conditioning. Specifically, we investigated whether changes in representation of GS, relative to CS+, stimuli over the course of the task related to individual tendency to generalize value from CS+ to GS stimuli – as captured behaviourally in avoidance responses on GS trials. We calculated a robust, cross-validated estimate of representational distance, Fisher’s linear discriminant contrast (see Methods, **Figure 4a**) in order to maximise the reliability of our results. Importantly, the use of a cross-validated distance measure means that derived (dis)-similarity estimates are unbiased by noise, and therefore have a meaningful zero point (Walther et al., 2015).

**Figure 4:**
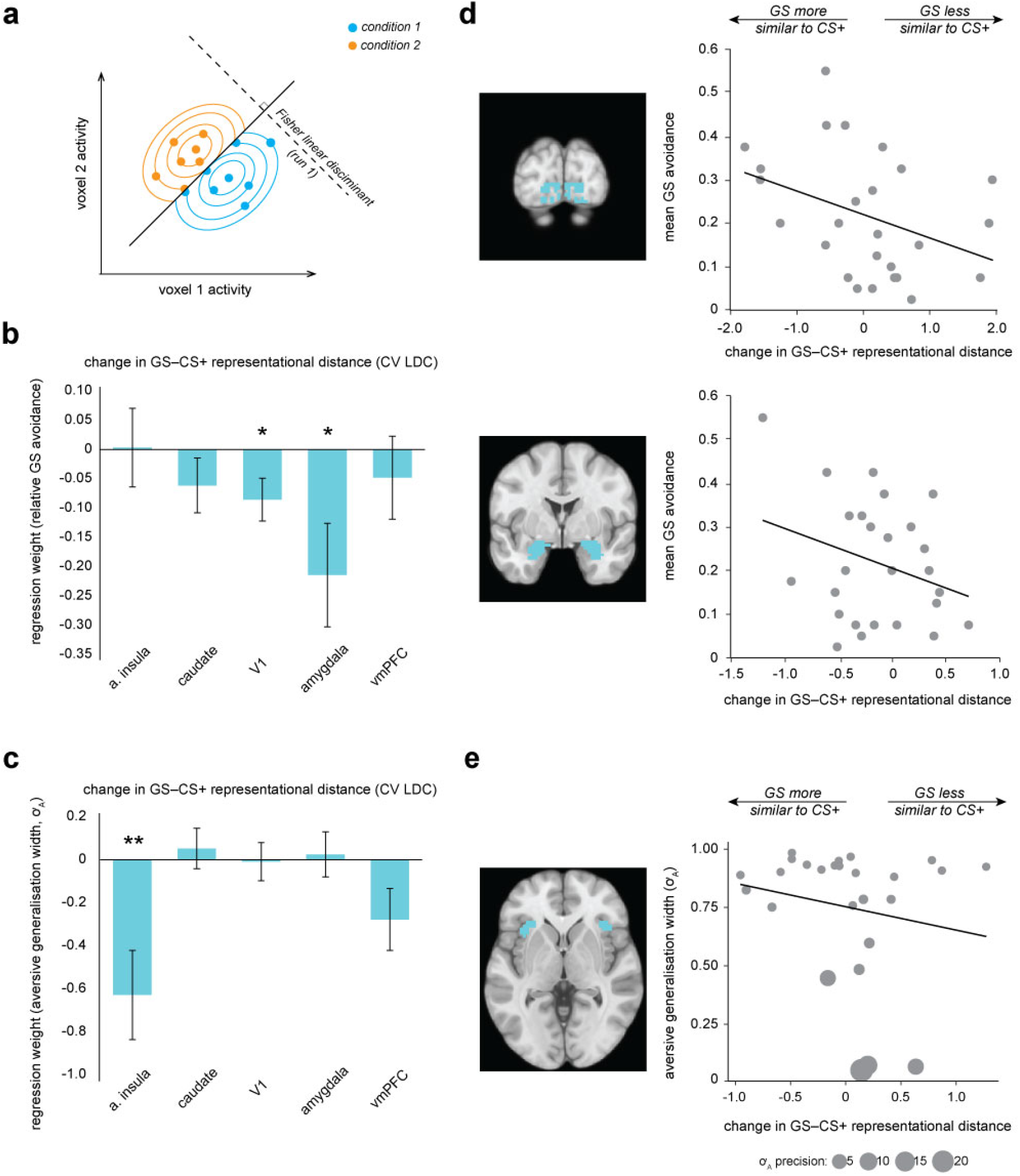
Multivariate fMRI results highlight regions where change in representational geometry over the course of the task between generalization stimuli (GSs) and pain-associated stimuli (CS+s) is related to individual differences in overall GS avoidance and the model parameter governing width of generalization from aversive feedback (σ_A_). **a**, Schematic of linear discriminant contrast analysis for a single imaging run (adapted from Kriegeskorte et al., 2007). **b,** Regression models detailing how changes in representational (dis)similarity earlier (blocks 1-2) vs later (blocks 3-5) over the course of the task in each ROI relate to relative proportionate avoidance on generalization trials, and c, in individual differences in model parameter governing width of aversive generalization. Error bars represent standard error. **d,** Visualisation of ROI volumes (pale blue shading) and bivariate relationships between change in representational geometry and raw GS avoidance in primary visual cortex and amygdala, and **e,** with individual σ_A_ values (in the anterior insula) weighted by individual parameter estimate precision (1/posterior variance). Larger bubble size represents greater precision (and therefore higher regression weight). CV LDC, leave-one-out cross-validated linear discriminant contrast; a insula, anterior insula; vmPFC, ventromedial prefontal cortex, \**p*<0.05, \*\**p*=0.007

Overall, for no region of interest was there a significant group level change in representational distance between GS and CS+ stimuli (there was a non-significant trend for GSs to come to be more similarly represented to the CS+ stimuli in the amygdala *p*=0.146, all other *p* values >0.2, paired-sample *t* tests). However, across individuals, greater increase in similarity of representation of GS to CS+ stimuli over the course of the task in both V1 (*p*=0.0288) and the amygdala (*p*=0.0245) was related to greater behavioural generalization in terms of greater relative GS avoidance (multiple linear regression model; **Table 1a, Figure 4b**). For individuals who made a higher relative proportion of avoidance responses towards generalization stimuli, V1 and amgydalar representations of GS stimuli came to be more similar to those of CS+ stimuli over the course of the task – but for individuals who avoided less on GS trials, GS stimuli came to be less similarly represented to CS+s in these regions (for visualisation of relationships between raw proportionate GS avoidance and distance measures, see **Figure 4d**). There was no evidence of a significant relationship between GS-CS+ representational distance change and relative GS avoidance in the anterior insula, striatum, or vmPFC **(Table 1a, Figure 4b)**. We confirmed these results by implementing a cross-validated regularised regression (CV LASSO, see Methods) on the same data (this kind of regression shrinks non-significant predictor coefficients to zero, and generally results in smaller coefficients compared to traditional linear regression). Under this robust approach, changes in GS-CS+ similarity in both V1 and the amygdala, but not other regions, were retained in the model as significant predictors of relative GS avoidance (*β*=-0.030, -0.059, respectively) in the model that minimised mean squared error (MSE).

**Table 1a.** Changes in representational distance (cross-validated LDC) with conditioning: relationship to overall generalization stimulus (GS) avoidance.

| Change in GS –<br>CS+<br>representational<br>distance | $\beta$ | SE | $t$ | $p$ |
| --- | --- | --- | --- | --- |
| a. insula | 0.0032 | 0.0672 | 0.048 | 0.962 |
| caudate | -0.0614 | 0.0466 | -1.318 | 0.202 |
| amygdala | -0.214 | 0.0881 | -2.432 | 0.025* |
| V1 | -0.0857 | 0.0364 | -2.355 | 0.029* |
| vmPFC | -0.0482 | 0.0709 | -0.679 | 0.505 |

Using a *post hoc* test, we examined whether changes in GS-CS+ representational distance in V1 might relate to changes in absolute discriminability of generalization stimuli (as measured on the day before and day after the generalization test session). Mean discriminability for GSs (CS+ ± θ) was 0.75 on session 1, by definition, and 0.79 on session 3 (± 0.14, range 0.465–0.994; although note at the group level there was no significant change in θ values measured across sessions, see above). Under this exploratory analysis, we found evidence of a significant association between change in V1 GS-CS+ representational distance during the task, and postconditioning changes in perceptual discriminability of the GSs. Individuals who showed an increase in similarity of representation showed worse perceptual performance post-(vs pre-) conditioning, and those who showed decreased similarity showing better performance (Spearman’s ρ=0.504, *p*=0.009; see **Figure S4)**. There was no significant relationship between change in perceptual acuity and representational distance in any other brain region (all p>0.010, Bonferroni corrected p value for multiple independent tests, alpha=0.05).

All the univariate fMRI findings presented above remained significant if re-ran using regressors derived from a model where perceptual discriminability of GSs changes linearly over the course of the task from pre- to post-conditioning measured acuity levels (see Supplementary Results, full statistical maps for all analyses available at Neurovault; neurovault.org/collections/3177).

## Changes in neural representation of generalization stimuli over the course of the task and relationship to individual differences in value-based generalization

We also sought to relate individual changes in similarity of representation of GS towards CS+ stimuli over the course of the to task to individual model parameter estimates governing width of generalization, specifically from aversive feedback (σ_A_ values).

We found that greater increases in similarity of representation of the GS relative to CS+ stimuli over the course of the task in the anterior insula were related to larger generalization from aversive feedback parameter estimates (*p*=0.0065, all other regions *p*>0.06; precision-weighted multiple linear regression model; see **Table 1b, Figure 4c,e**). However, this result did not survive in the more robust CV LASSO model, where representational change in no region was retained as a significant predictor.

**Table 1b.**
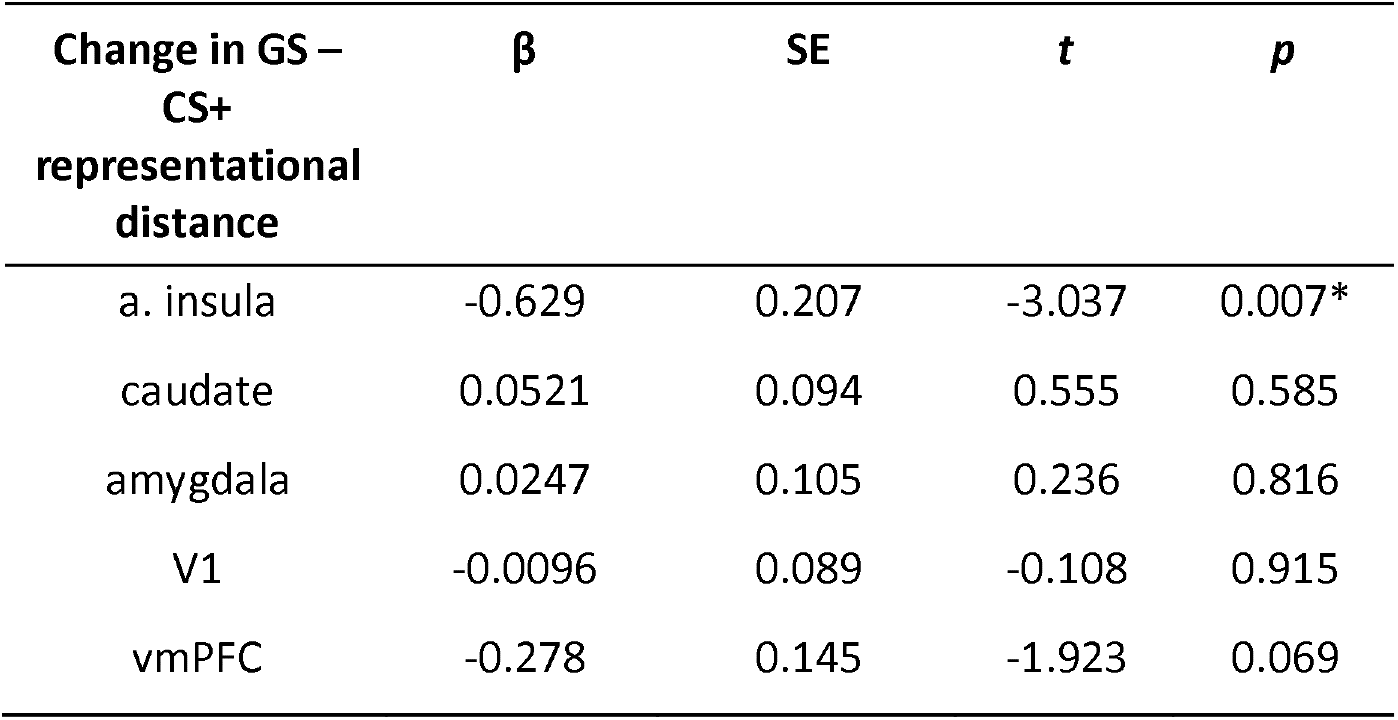
Changes in representational distance (cross-validated LDC) with conditioning: relationship to model parameter governing width of generalization from aversive feedback (σ_A_). A. insula, anterior insula; vmPFC, ventromedial prefrontal cortex; V1, primary visual cortex; SE, standard error. \**p*≤0.05

We hypothesised that the reason that representational change in the anterior insula, but not amygdala, might be identified as relating to aversion-specific aspects of avoidance generalization is that the amygdala may also play a role in generalization about neutral (or safe) outcomes. Although less well-studied compared to the aversive domain, there is evidence that the amygdala is involved in the acquisition of information about safety signals in rodents and non-human primates (Rogan et al., 2005; Genud-Gabai et al., 2013), and that medial prefrontal entrainment of the amygdala is associated with learned safety (successful overcoming of generalized conditioned fear) in mice (Likhtik et al., 2014). This fits with a large literature on the vmPFC playing a role in ‘safety signalling’ in humans (Fullana et al., 2016).

As a further exploratory analysis, we therefore investigated whether there was a relationship between change in GS-CS- similarity over the course of the task in the amygdala and vmPFC and individual values of the parameter governing width of generalization from neutral (non-pain) feedback, σ_N_. (*Nb*, due to the arrangement of task stimuli, see **Figure 1b,** our design is not optimised to probe GS-CS- value generalization at the stimulus category level.) We found evidence of a significant relationship between GS-CS- similarity change in the vmPFC and individual σ_N_ values -such that individuals where representation of GSs came to be more similar to CS- in this region had greater neutral (‘safety’) generalization parameter values. There was no strong evidence for a similar relationship in the amygdala (vmPFC: β=-0.100, SE 0.04, *t*=-2.379, *p*=0.026; amygdala: *β*=-0.037, SE 0.03, *t*=-1.296, *p*=0.207; precision-weighted multiple linear regression model). Neither predictor was retained in the CV LASSO model.

## Relationship between individual differences in value-based generalization and selfreported psychopathology

Hypotheses about the role of generalization in psychological disorders tend to relate to an over-generalization of aversive information – but it has also been proposed that poor discrimination (e.g. between CS+ and CS- in anxiety groups) may be due to inadequate learning about safety cues (see Introduction). We therefore looked first at how psychological symptoms scores related to individual σ_A_ values, but also examined possible relationships with individual σ_N_ values, in our online cohort (*N*=482).

Following the approach of Gillan and colleagues (Gillan et al., 2016), the online group completed a battery of self-report questionnaires that probed symptoms hypothesized to be related to aversive over-generalization (trait anxiety, mood disorder symptoms, obsessive-compulsive traits, and ‘global’ cognitive style), in addition to some positive control measures (apathy and impulsivity scales). (A summary of scores on these measures and other demographic information for both samples is available in **Table S2**.) To enable comparison with the findings of Gillan et al., self-report information was first compared to individual parameter estimates using precision-weighted linear regression models, controlling for age and gender identity (see Methods). This approach was then complemented by the implementation of cross-validated regularised regression models (CV LASSO regression), as in the previous section (these models also included age and gender information as regressors of no interest).

First, we sought to identify whether individual values of the parameter governing width of generalization from aversive feedback (σ_A_) were related to symptom scores on any measure. Total scores across measures exhibited good to excellent internal reliability (mean Cronbach’s α=0.882, see **Table S3**), and, as might be expected, covaried significantly across participants (mean absolute r for inter-correlation between scores=0.479). Regression of total scores against parameter estimates was therefore implemented in separate models for each measure, in order to enable meaningful partition of variance (as per Gillan et al., 2016). The Nyholt-Bonferroni corrected p value for significance across these separate models of non-independent measures was *p*<0.010 to maintain an alpha of 0.05 (effective number of independent variables=5.0, see Methods).

Parameter estimates governing width of generalization from aversive feedback were found to be significantly positively associated with trait anxiety scores (greater width with greater anxiety), and significantly negatively associated with trait apathy (smaller width with greater apathy; anxiety, *p*=0.009, apathy, *p*<0.001, individual precision-weighted linear regression models controlling for age and gender; see **Table 2a, Figure 5a**). These two effects remained significant when trait anxiety and apathy scores were included in the same model, suggesting they were independent (anxiety: β=0.050, SE 0.015, *t*=3.34, apathy: β=-0.060, SE 0.014, *t*=-4.28; both *p*<0.001). This result was confirmed under the cross-validated and regularised analysis; when all predictors were entered in the same model both anxiety and apathy total scores were retained as predictors in the model that minimised MSE (β=0.021, β=-0.032, respectively). No questionnaire total scores were significantly related to σ_N_ values (*p*>0.05).

**Figure 5:**
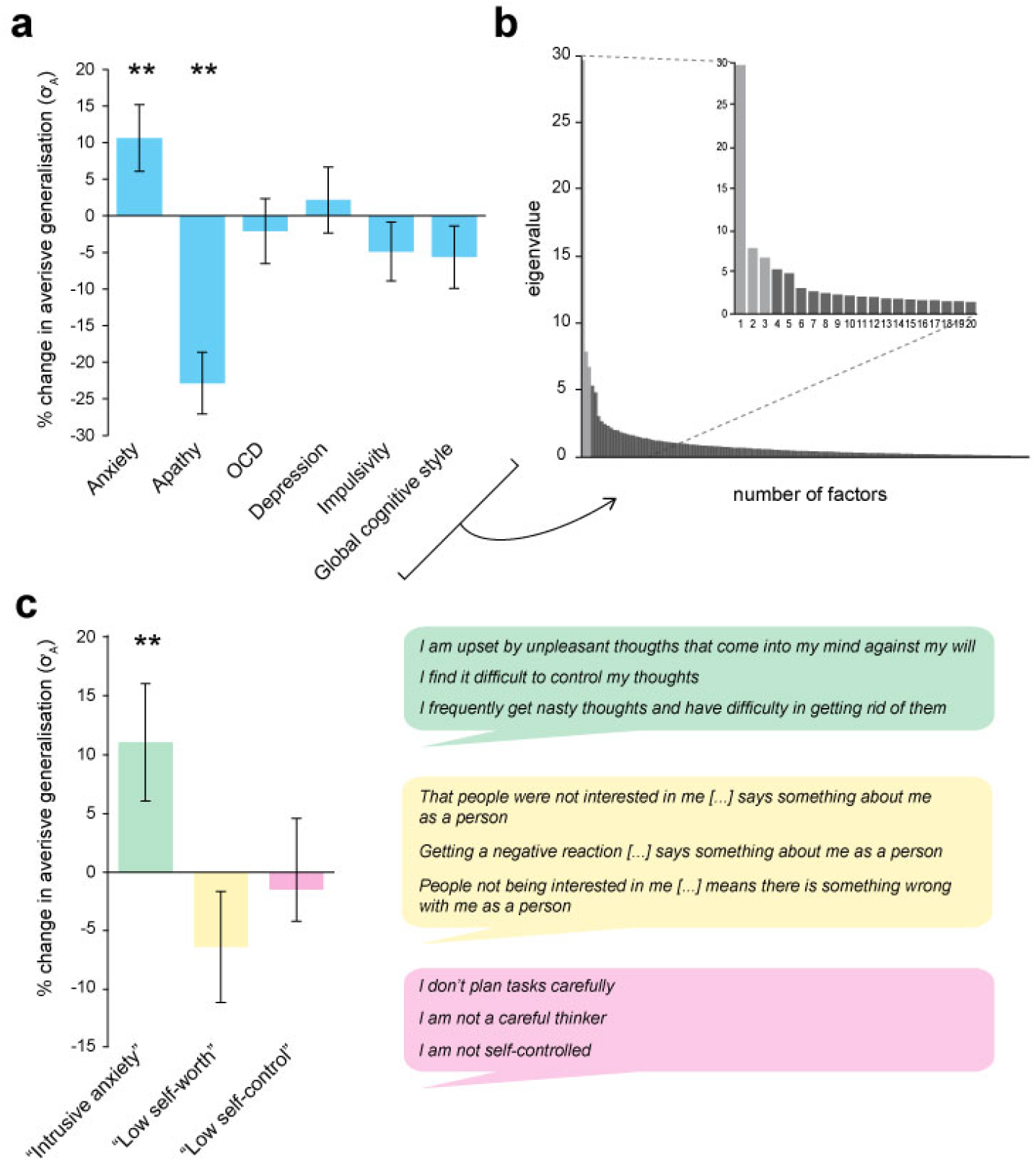
Associations between individual differences in aversive generalization and psychological symptom scores. **a**, Percentage change in the model parameter governing width of generalization from aversive feedback (σ_A_) with a 1 standard deviation increase in total score on each individual questionnaire measure used (individual regression models), **b,** Scree plot indicating results of a factor analysis in which all response items from these measures (*N*=142) were entered (inset, first 20 factors). A three-factor solution (lighter shaded bars) was indicated as the most parsimonious structure. **c,** Percentage change in σ_A_ with an increase in 1 SD for each of the factor analysis-derived symptom scores (single regression model). The right hand panel shows the top three loading items for each factor, which were used to derive factor labels. Error bars represent standard error. \*\**p*<0.009

**Table 2a.** Relationship between width of generalization from aversive feedback (σ_A_ value estimates) and questionnaire total scores. Each line represents the results of a separate model, as questionnaire scores were significantly collinear. STAI, Spielberger State-Trait Anxiety Inventory (trait scale); AMI, Apathy Motivation Index; OCI-R, Obsessive-Compulsive Index (Revised); PHQ9, Physician’s Health Questionnaire 9 (a brief measure of mood disorder symptoms); BIS-11, Barratt Impulsivity Scale (version 11); CSQ global, Cognitive Style Questionnaire cognitive globalisation score. SE, standard error. \**p*<0.010 (Nyholt-Bonferroni corrected *p* value for multiple tests on non-independent data, alpha=0.05).

| Questionnaire measure | $\beta$ | SE | $t$ | $p$ |
| --- | --- | --- | --- | --- |
| STAI total | 0.039 | 0.015 | 2.626 | 0.009* |
| AMI total | -0.051 | 0.014 | -3.687 | <0.001* |
| OCI-R total | 0.005 | 0.014 | 0.373 | 0.710 |
| PHQ9 total | 0.021 | 0.015 | 1.476 | 0.141 |
| BIS-11 total | -0.005 | 0.013 | -0.410 | 0.682 |
| CSQ global | -0.014 | 0.014 | -0.978 | 0.328 |

**Table 2b.** Relationship between generalization width from aversive feedback (σ_A_ value estimates) and factor analysis-derived symptom scores. All factor scores were included in the same model. SE, standard error. \**p*<0.05

| Factor analysis-derived symptom score | $\beta$ | SE | $t$ | $p$ |
| --- | --- | --- | --- | --- |
| "Intrusive-anxiety" | 0.043 | 0.016 | 2.677 | 0.008* |
| "Low self-worth" | -0.019 | 0.015 | -1.255 | 0.210 |
| "Lack of self-control" | -0.000 | 0.014 | -0.032 | 0.975 |

As per Gillan et al, we also sought to reduce collinearity in our battery of self-report measures by entering all recorded items (*N*=142) into a factor analysis. Using an identical method to that described in the previously cited paper (see Methods), we derived a three-factor solution (for scree plot see **Figure 5b)**. These factors were labelled “intrusive anxiety”, “low self-worth”, and “low self-control” on the basis of their top loading items (see **Figure 5c**).

The “intrusive anxiety” factor was mostly composed of items from the trait scale of State-Trait Anxiety Inventory (STAI; 20 items, mean loading=0.457 ±0.12), Obsessive-Compulsive Index (OCI; 18 items, mainly items probing intrusive thoughts and checking behaviour, mean loading=0.602 ±0.087), Physician’s Health Questionnaire (PHQ9; 8 items probing mood disorder symptoms, mean loading =0.531 ±0.056), and the Barratt Impulsivity Scale (BIS; 6 items pertaining to racing/intrusive thoughts and restlessness, mean loading=0.386 ±0.15). “Low self-worth” was mostly comprised of items from the Cognitive Style Questionnaire (CSQ; 37 items, mainly from low self-worth and internal attribution subscales, mean loading=0.518 ±0.13) and the STAI (11 items, mainly related to low self-worth/negative self-affect, mean loading=0.322 ±0.054). “Low self-control “mostly comprised items from the BIS (23 items, mainly from the non-planning and attentional impulsivity subscales, mean loading=0.485 ±0.15), with some loading from the apathy motivation index (AMI; 6 items from the behavioural amotivation subscale, mean loading=0.356 ±0.093) and STAI (7 items relating to feel uncontent/unrested, mean loading 0.321 ±0.04). (For full item loadings for each factor, see **Table S3**.)

The “intrusive anxiety” factor analysis-derived symptom score was significantly and selectively related to individual differences in aversive generalization width (σ_A_ values) – in both multiple linear and robust regression models (p=0.008, precision-weighted multiple regression model; see **Table 2b, Figure 5c;** only factor retained in MSE-minimising CV LASSO model, *β*=0.019). None of the factor analysis-derived symptom scores were related to individual σ_N_ values (all *p*>0.1).

## Discussion

The results presented here provide robust evidence for generalization in human avoidance learning. In particular, we demonstrate that generalization involves a number of distinct processes relating to different components of avoidance: perceptual uncertainty, aversive value generalization, and neutral (safety) value generalization. These processes each relate to different patterns of neural representations in the brain. Finally, we show that aversive value generalization is a specific predictor of trait anxiety in a large population sample.

Examining instrumental avoidance behaviour allows us to investigate how individuals learn about and attribute value to the set of *actions* they can take when faced with a particular stimulus or situation (as distinct from passively learnt Pavlovian stimulus-value associations). Using reinforcement learning modelling, we found behavioural evidence for additional value-based contributions to avoidance generalization (i.e. over and above that which might be expected from perceptual uncertainty alone) in two independent groups of participants (sampling different populations, and using two different kinds of aversive reinforcer). Notably, choice data from both groups supported an account of value-generalization that allowed for different widths of generalization from aversive (pain or monetary loss) *vs* neutral (no pain or loss) feedback. Consistent with previous evidence from studies of generalization of Pavlovian conditioning in humans and non-human primates, we observed larger width generalization functions for aversive compared to neutral feedback (Schechtman et al., 2010; Resnik and Paz, 2015; Laufer et al., 2016). In both groups, estimates of free parameters governing widths of these two processes were uncorrelated, suggesting they might relate to at least partially separable mechanisms.

This model-based approach enabled us to identify brain regions where BOLD signal was related to variance in modelled quantities specific to value-based generalization (specifically, expected value and prediction error signals). Both the anterior insula and the caudate have been previously identified in representing perceptual-similarity derived generalization gradients following Pavlovian fear conditioning in humans (Dunsmoor et al., 2011; Greenberg et al., 2013; Lissek et al., 2014). Here, we provide evidence that BOLD signal in these regions is specifically related to additional value-based generalization processes, when potential perceptual confusion between visually similar task stimuli has been accounted for.

The anterior insula and striatum (more ventrally) have previously been implicated in representing expected value and prediction error signals in higher-order pain conditioning (Seymour et al., 2004), and the dorsal striatum is implicated in prediction error signals in avoidance learning (Palminteri et al., 2012; Seymour et al., 2012; Eldar et al., 2016), suggesting an important role for these structures in aversive learning (see also Delgado et al., 2009). Dorsal, rather than more ventral striatal control has also been implicated in the transfer from goal-directed to habit-based avoidance in instrumental paradigms (LeDoux et al., 2017). Greater understanding of habitual control in excessive avoidance has particular clinical relevance as it may explain why maladaptive avoidance can persist following extinction (e.g. contributing to treatment-resistance in exposure therapy for anxiety disorders; Treanor and Barry, 2017), and has been proposed as core mechanism in obsessive-compulsive disorder (Gillan et al., 2014). It is important to note however that in the current study, our dorsal striatal findings did not reach significance for the prediction error analysis (at least in the constant perceptual acuity model), and we may wish to treat significant findings from the expected value analysis in this brain region with some caution due to potential partial confounding with motor preparation responses (see Methods). We found no evidence of univariate encoding specific to value-based model quantities in the amygdala, primary visual cortex (V1), or ventromedial prefrontal cortex (vmPFC). However, this may be because this kind of analysis is not ideally suited to detect distributed representations involved in associative learning.

In previous studies of Pavlovian aversive conditioning, it has been demonstrated that positively conditioned stimuli come to be more closely represented to the primary aversive outcome in multivariate space (e.g. across neural ensemble activity in the basolateral amygdala, Grewe et al., 2017). Here, we used a robust, cross-validated measure of representational distance to analyse data across all voxels in regions of interest, and found that increased similarity of representation of GS to CS+ stimuli over the course of the task in both the amygdala and primary sensory cortex was related to higher overall behavioural generalization (higher proportionate avoidance on generalization trials). Individuals for whom GS stimuli came to be more closely represented to CS+s in these brain regions (despite never having been directly associated with the aversive outcome) chose to avoid more in the face of GS stimuli – and *vice versa*. This change in representational geometry, in association with the lack of opportunity for extinction of inappropriately generalized value in an avoidance context, may have contributed towards the stability of generalization (in terms of overall GS avoidance) we observed over the later phases of the task.

Consistent with perceptual accounts of generalization, a post-hoc analysis suggested that representational change for GSs relative to CS+ stimuli over the course of the task in primary visual cortex might account for some of the generalization in avoidance we observed (in addition or parallel to value-based mechanisms identified above). Individuals who avoided more frequently on generalization trials, and who showed associated increases in GS-CS+ representational similarity in V1, exhibited decreased perceptual acuity for task stimuli on next day perceptual testing – with the opposite pattern observed in participants who showed lower GS avoidance. Absolute decreases in discriminability for task stimuli result in increased generalization ‘for free’ (without having to involve additional mechanisms), and therefore may contribute to maintenance of generalization in some participants.

However, consistent with accounts that favour the involvement of a wider network of brain regions in coordinating generalization across stimuli, we also found a role for multivariate anterior insula representations specifically in individual differences in aversive value generalization. Individuals who had higher estimates for the model parameter governing value generalization specifically from aversive feedback showed greater increases in similarity of GS-CS+ representation in the anterior insula, but not other regions of interest (e.g. notably the same relationship was not evident in the amygdala). Although less well optimised, we conducted a further analysis to probe whether changes in GS relative to CS- stimuli might be associated with individual estimates of the model parameter governing width of generalization specifically from neutral (or ‘safe’) outcomes (in this case, omission of painful shock). We restricted this exploratory analysis to the amygdala and vmPFC, on the basis of previous evidence that these regions may play a role in the generalization of information regarding safety or aversive outcome omission (Likhtik et al., 2014; Fullana et al., 2016). Although we found no strong evidence for a relationship between safety generalization and GS-CS- representation in the amygdala, increased representational similarity of GS to CS- stimuli over the course of the task in the vmPFC was associated with higher neutral generalization parameter values.

Although the vmPFC has previously been demonstrated to show inverse perceptual similarity-derived generalization gradients following aversive conditioning (e.g. Lissek et al., 2014; Onat and Büchel, 2015), it is not always clear from the experimental design whether this represents the simple inverse of aversive gradients (stemming from the CS+), or rather the positive signalling of safety gradients (stemming from the CS-). The evidence presented here provides tentative support for the latter account, at least in an instrumental context.

Excessive avoidance in response to contexts or stimuli which do not pose a threat to an individual’s health or well-being can significantly impair general functioning and is often associated with high levels of psychological distress (Arnaudova et al., 2017). Such maladaptive avoidance has been identified as a core pathological dimension across several psychological disorders, including anxiety disorders, obsessive-compulsive disorder, chronic pain, and depression (LeDoux et al., 2017). Over-generalization of aversive feedback to encompass non-threatening but psychologically similar stimuli or contexts has been proposed as a key mechanism underlying the initiation and maintenance of excessive avoidance in these conditions (Duits et al., 2015; Dymond et al., 2015; Harvie et al., 2017; Pearson et al., 2015) – however the link between generalization of negative value and inappropriate avoidance behaviour has been relatively underexplored.

We found selective relationships between psychological symptom scores and individual parameter estimates governing width of value generalization from aversive, but not safe/neutral outcomes. The largest positive relationship between symptom score and magnitude of aversive generalization was for the factor-analysis derived score labelled “intrusive anxiety”, which mainly comprised items probing self-reported trait anxiety, but also reports of intrusive thoughts from the obsessive-compulsive inventory (% increase in parameter value with a 1SD increase in symptom score was 11.0% for intrusive anxiety, and 10.6% for trait anxiety alone). We also found a significant negative relationship between self-reported apathy and aversive generalization (22.9% decrease in parameter value with a 1SD increase in symptom score) – an effect which appeared to be independent from that relating to self-reported anxiety. This is an interesting finding, as we often think about apathy as involving a greater sensitivity to perceived effort, or decreased sensitivity to potential rewards, rather than a decreased impact of information about punishments (e.g. Bonnelle et al., 2015). As, to our knowledge, there has been no previously reported link between self-reported apathy and aversive generalization, this finding would benefit from future replication.

In summary, the findings reported here demonstrate the benefits of parsing complex processes such as generalization into separate components, and examining individual relationships between these components and both neural mechanisms and self-reported psychopathology. This approach may help unify previous apparently contradictory observations, and underlines that both perceptual and value-based processes are likely at work in generalization phenomena. Identification of patients across diagnostic categories who may have a primary deficit in excessive aversive generalization may help target them towards treatments which work more effectively. Further, greater understanding of the mechanisms of over-generalization of avoidance (e.g. transfer to habit-based control systems) may help improve understanding of treatment resistance in these disorders.

## Author Contributions

AN. TWR, and BS designed the study. AN collected the data and conducted the analysis. AN, TWR, and BS wrote the paper.

## Acknowledgements

This study was funded by the Wellcome Trust (grant number 097490/Z/11/A to BS). TWR was funded by a a Wellcome Trust Senior Investigator Award (grant number 104631/Z/14/Z). The authors declare no relevant conflicts of interest.

## Methods and Materials

### Code and data availability

All relevant code for stimulus generation, data collection, and data analysis, in addition to raw behavioural data, is available at the project’s Open Science Framework page (osf.io/25t3f). Raw functional imaging data is deposited at openfMRI *(url to follow)* and derived statistical maps are available at NeuroVault (neurovault.org/collections/3177).

## Design

### fMRI sample

#### Protocol

Each participant completed three testing sessions on three consecutive days. On the first day, participants were pre-screened, gave informed consent, and performed initial sensory acuity testing for the generalization task stimuli. On the second day, participants completed the generalization of instrumental avoidance task (performed in fMRI scanner, using individually-generated conditioning stimuli [CSs] derived from day one perceptual performance), followed by visual analogue scale (VAS) ratings of pain expectancy for each CS. On the third day, participants repeated the perceptual acuity test.

All participants were recruited via online advertisement. Exclusion criteria were left-handedness and history of neurological or psychological illness, in addition to usual MR safety criteria. The sample size was chosen on the basis of a power calculation. Previous functional imaging studies in humans have found effect sizes in the region of r=~0.5 for generalization-related BOLD signal and individual difference measures (Greenberg et al., 2013; Cha et al., 2014; Lissek et al., 2014). We calculated that a sample of *N*=26 would allow us to detect r=0.5 with an alpha of 0.05 and power of 80%, two-tailed. Volunteers were paid £20 per hour in recompense for their time and discomfort arising from the painful electrical stimulation. The study was approved by the University of Cambridge Psychology Research Ethics Committee.

#### Delayed-punished perceptual discrimination task

Prior to starting the task, participants were introduced to the shock and electrode and a work-up procedure was performed (as described below) to set the level of painful stimulation. The delayed-punished perceptual task was then carried out, as summarized in **Figure 1b**. Briefly, on each trial, participants viewed an individual shape (target or comparison stimulus, order randomized on each trial), followed by a mask (scrambled mean shape image), delay period (blank screen), second shape, and second mask. At the end of each trial, participants had to indicate whether they thought the two shapes had been the same, or different. The inter-stimulus delay period of four seconds was chosen to be long enough such that comparison of stimuli could not be achieved by instantaneous mechanisms, but required comparison in short-term memory (e.g. primate data suggests discrimination performance for visual features decreases significantly from <1s to around 4–5s inter-stimulus delay; Pasternak and Greenlee, 2005), and roughly matched to the inter-trial interval from the generalization task. There were 16 trials per absolute value interval per target (160 trials total), and trials were divided into 4 equal blocks. At the end of each block, participants received feedback on how many incorrect judgments they had made, and received a proportionate number of painful electric shocks as punishment (roughly one painful shock per five incorrect judgments).

Stimuli were 5-fold radially symmetric flower-like shapes, as described in van Dam and Ernst, 2015. These were selected on the basis of previous psychophysical testing demonstrating that they are perceptually linear, and the fact that they can be individually generated along axis a single perceptual axis of ‘spikiness’ using the mathematical description provided in the paper. Shape ‘spikiness’ is parameterized by a single value, *ρ* (where 0 < *ρ* < 1), which relates the inner and outer radii of the shape such that stimuli are of constant surface area. Target stimuli were shapes with ρ values of the two CS+ stimuli from the generalization task (0.25 and 0.75). These target stimuli were compared to comparison stimuli of intervals of ± 0, 0.05, 0.075,0.1, and 0.15 *ρ*, such that the possible range of different shapes was well tiled. Participants worked on a pre-defined set of comparison stimuli (opposed to a staircased approach) so that pre-exposure to conditioning task stimuli (and therefore opportunity for perceptual learning) would be matched across individuals.

#### Generalization of instrumental avoidance task (pain version)

Participants completed five blocks of 38 trials each. On each trial, participants were presented with a stimulus in the centre of their screen. This initiated a 3s decision period, during which they must decide whether or not to make an ‘escape’ (avoidance) response. Following this, a yellow bounding box appeared around the shape, indicating the time when an avoidance response could be made was over and they would receive the outcome for that trial. If an avoidance response was made, no shock was ever delivered on that trial. If no avoidance was made, and the stimulus was a ‘safe’ shape (CS-), no shock was delivered. If the stimulus was a ‘dangerous’ shape (CS+), a painful shock was delivered on 80% of non-avoidance trials at the end of this outcome period (i.e. 6s from stimulus onset, see **Figure 5a)**. On a low frequency of trials, shapes were generalization stimuli (GSs). These stimuli were individually generated to be 75% reliably distinguishable from adjacent CS+s based on day one perceptual task performance (see **Figure 1b),** and were never associated with painful shock.

The stimulus array was asymmetric in perceptual space (see **Figure 1b),** with two CS+ (and four associated GS) stimuli – one nearer and further from an intermediary CS-. This array was chosen in order to probe the presence of characteristic asymmetries in conditioned responses that are hypothesised to arise from the interaction of oppositely signed generalization gradients (e.g. peak shift, Hanson, 1959), and on the basis of previous observations that change in perceptual discriminability of aversively conditioned stimuli (CS+s) may depend on the relative ‘nearness’ of safety stimuli (CS- s) in perceptual space (Aizenberg and Geffen, 2013). Axis direction (in terms of increasing or decreasing ‘spikiness’) was counterbalanced across participants. Trial types were presented in the following ratio: 10 CS- : 10 CS+(*2): 2 GS in a pseudorandom sequence, in order to minimise learning about GS stimuli.

### Online sample

#### Protocol

In order to test relationship with real-world psychological symptoms in an appropriately powered sample, an online version of the study was also carried out, following the approach of Gillan et al. (Gillan and Daw, 2016; Gillan et al., 2016). Participants were Amazon Mechanical Turk (AMT) workers based in the USA (in practice, had an AMT account linked to US bank with provision of an US social security number). Participants were required to be over 18 years of age, but otherwise remained anonymous.

Participants completed an online consent procedure, and provided limited demographic information (age and gender identity). They then read several screens of detailed task instructions (including visual examples of sample trials), based on the standardized instructions given to lab study participants. Participants were required to pass a 10 item true/false quiz on task structure before continuing (scoring less than 10/10 returned participants to the instruction screens). They then performed a monetary loss-based version of the generalization of instrumental avoidance task (see below), followed by a battery of questionnaires probing psychological symptoms and cognitive style.

We calculated that a final sample size >459 should be powered to detect a small effect size of 0.15 (association between behavioural and self-report parameters), at 80% power (two-tailed). As expected attrition following quality control was ~15% (Gillan et al., 2016), we collected *N*=550 complete datasets, yielding a final expected sample size of ~468.

Payment rates were based on UK ethical standards for online experiments (equivalent to a minimum of £5ph). Participants were paid a flat rate of $2.50 for taking part, plus up to around $3.00 additional bonus payment depending on task performance. The average bonus payment was $2.21 (± 0.82) and the average time between accepting and submitting the task was 42 minutes (equivalent to $6.72 mean hourly payment rate). The study was approved by the University of Cambridge Psychology Research Ethics Committee.

#### Generalization of instrumental avoidance task (loss version)

The generalization task was identical in structure to that performed by the lab-based participants, but used monetary loss instead of painful shock as the aversive reinforcer **(Figure 1c)**. Prior to starting the task, participants were endowed with a $6.00 stake, and instructed that, although a certain amount of loss was inevitable, whatever total remained at the end of the task would be paid directly to them as a bonus (the loss therefore had real-world value). As BOLD data was not being collected, trials were slightly shorter than for the fMRI group (second set of timing figures, **Figure 1c)** – although the length of the decision period was kept the same.

Perceptual testing was not performed in the online sample due to time constraints, and the inability to control the testing environment (e.g. participant distance from screen, window size, etc.) over the course of testing. Generalization stimuli were therefore the same for all participants, and generated on the basis of mean perceptual performance on the delayed-punished perceptual task in a pilot sample.

#### Questionnaire battery

Following completion of the generalization task, participants completed several self-report measures (questionnaire order was randomised across participants). These measures were chosen to probe psychological constructs hypothesized to be related to over-generalization of aversive outcomes (anxiety, depression, and obsessive-compulsive symptomatology), as well as positive controls that might suggest a more general effect of psychopathology on task performance (impulsivity, apathy). Questionnaires consisted of the trait scale of the State-Trait Anxiety Inventory (STAI; Spielberger et al., 1970); the Physician’s Healthy Questionnaire 9 (PHQ9; Martin et al., 2006), a brief measure of mood disorder symptoms; the revised (short-form) Obsessive-Compulsive Index (OCI-R; Foa et al., 2002); the Barratt Impulsiveness Scale v11 (BIS-11; Patton et al., 1995); and the Apathy Motivation Index (AMI; Ang et al., 2017). All chosen measures have previously been shown to be suitable for use in the general population.

A short version of the Cognitive Style Questionnaire (CSQ-SF; Meins et al., 2012) was also administered. This self-report measure asks participants to imagine themselves in various scenarios (e.g. “Imagine you go to a party and people are not interested in you”), and then probes the imagined causes of this scenario along dimensions of “internal”, “global”, and “stable” attributions, plus low self-worth. On this measure, a more “global” cognitive style reflects a tendency to attribute negative events to causes which are general, rather than specific (a cognitive form of over-generalization), and has been found to be a predictor of future depressed mood (Pearson et al., 2015). The CSQ-SF was administered at the end of the battery of questionnaires for all participants in order to avoid possible mood-induction effects.

#### Quality control procedure

Following previous studies utilizing AMT (Crump et al., 2013; Gillan et al., 2016), a number of exclusion criteria were applied sequentially to the dataset to attempt to exclude poor quality responses. Firstly, we excluded participants who made avoidance responses on less than 50% of total CS+ trials (indicating lack of learning/random responding on these trials), *N*=62. Secondly, we further excluded participants who selected the wrong answer to a catch item inserted into the questionnaire battery (“Please select the answer “a little” if you are reading this question”), *N=*6. 68 datasets were excluded in total (12.3% of those collected), yielding a final sample size of 482. Questionnaire data quality was further assessed via calculation of internal reliability coefficients for each measure (Cronbach’s α).

## Data collection

### fMRI sample

Stimulus presentation and response collection was coded using Cogent2000 v1.30, run in Matlab R2015b (Mathworks). Perceptual testing on day one and three took place in a laboratory, and generalization testing in an fMRI scanner. Size of stimuli in terms of visual angle subtended were matched between lab and scanner environments in order to ensure ~constant discriminability.

For the painful stimulation, electric current was generated using DS7A constant current stimulator (Digitimer), delivered to a custom fMRI compatible annular electrode (which delivers a highly unpleasant, pin-prick like sensation), worn on the back of the participant’s dominant (right) hand. All participants underwent a standardised intensity work-up procedure at the start of each testing day, in order to match subjective pain levels across sessions to a level that was reported to be painful, but bearable (8 out of 10 on a VAS ranging from *0* [“no pain”] to *10* [“worst imaginable pain”]). The pain delivery setup was identical for lab-based and MR sessions.

Functional imaging data were collected on a 3T Siemens Magnetom Skyra (Siemens Healthcare), equipped with a 32-channel head coil. Respiration data was collected during functional scanning using a pneumatic breathing belt (BrainProducts), and choice (avoidance) data was recorded using an MR-compatible button box.

Field maps were acquired in order to correct for inhomogeneities in the static magnetic field (short TE=5.19ms, long TE=7.56ms, 32x3mm slices). Five functional sessions of 212 volumes were collected using a gradient echo planar imaging (EPI) sequence (TR=2000ms, TE=30ms, flip angle=90⍰, tilt=-30⍰, slices per volume=25, voxel size 3x3x3mm) (this included 3 dummy volumes, in addition to 3 discarded by the scanner). Limited field of view (constrained by equipment used for additional physiological data collection) was aligned to the base of brain and angled away from the orbits, such that there was full coverage of the occipital and temporal lobes, plus prefrontal cortex. A T1-weighted MPRAGE structural scan (voxel size 1x1x1mm) was also collected. Full metadata are available at openfMRI (*url to follow*).

### Online sample

The experiment was coded in javascript using jsPsych (de Leeuw, 2015; available at github.com/jspsych/jsPsych), and was deployed to Amazon Mechanical Turk via the psiTurk engine (Gureckis et al., 2016; available at github.com/NYUCCL/psiTurk). The experiment was hosted in the cloud using an Amazon Web Services EC2 instance. A more detailed description of this setup is available at osf.io/mjgtr. The task was not made available on mobile devices (phones or tablets) in an attempt to ensure minimum screen size.

## Analysis

### Perceptual acuity

For fMRI sample participants, psychometric functions (logistic function with free parameters governing slope, bias, and lapse, or stimulus-independent error, rate) were fitted to response data from the perceptual task using the psignifit toolbox v2.5.6 (available at bootstrap-software.org/psignifit), run in Matlab. This implements the constrained maximum-likelihood method of psychometric function fitting described in (Wichmann and Hill, 2001).

Individual psychometric functions were then used to calculate the different in ρ value necessary for the comparison stimulus to be distinguishable from the target on 75% of trials (henceforth, *ϑ*).

### Instrumental avoidance behaviour

Avoidance behaviour was modelled using a set of modified SARSA algorithms (Sutton and Barto, 1998). Each stimulus was modelled as a different state, with the value of executing each action (*avoid* or *notAvoid)* in each state {*V_s,a_*) updated after each trial (*t*) on the basis of a simple Rescorla-Wagner rule – i.e. on the basis of difference between the predicted value of that state-action pair, and the actual outcome of each trial (*R_t_*; coded as 0 for no shock/no loss and −1 for shock/monetary loss). Formally,

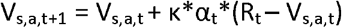

Learning rate (*α_t_*) was updated on each trial, according to the empirically well-supported Pearce-Hall associability rule (Pelley, 2004):

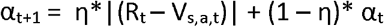

According to this rule, the learning rate on each trial is determined by the absolute magnitude of past prediction errors, such that state-action value estimates are updated by more when previous outcomes have been more surprising, and by less when they were less surprising. This allows for learning in terms of modelled value adjustment to be greater when outcomes are more surprising (e.g. at the start of the task), but to be lesser (leading to more stable values) when outcomes are better predicted. A non-constant learning rate also ensures that parameters governing width of value-based generalization, which are assumed to be constant over the course of the task, are identifiable during parameter estimation (see below equations). Individual differences in degree of dependence on prediction error history and overall scaling of learning rate are governed by the free parameters *κ* and *η*.

To model possible perceptual generalization / confusion between GSs and adjacent CS+s:

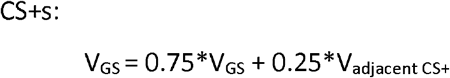

For the additional value-based generalization models, on each trial the values of all states were updated in proportion to their perceptual similarity to the current state, *i*, using a rule similar to those employed in previous studies (Kahnt et al., 2012; van Dam and Ernst, 2015) – i.e. according to a variable-width Gaussian function across perceptual space. For each state,; *j*:

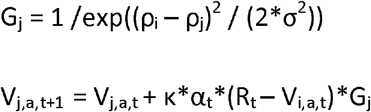

where *ρ* is the parameter governing shape ‘spikiness’, and the width of Gaussian function governing generalization is determined by the free parameter σ. For the fMRI sample, average *ρ* values were used for all subjects, as shapes had been matched in subjective perceptual space. For the 2-width model, different *σ* values were fit depending on whether the outcome for that trial was aversive or neutral (*σ_A_* and *σ_N_*, respectively).

As participants were explicitly instructed that they would never receive the aversive outcome if they made an avoidance response, the value of avoiding in any state (*V_s,notAvoid,t_*) was held constant at 0. Value estimates were fit to binary choice data via a softmax observation function, taking into account the cost of making an avoidance response (~additional 1/5 shock or unit monetary loss to be received at the end of that block):

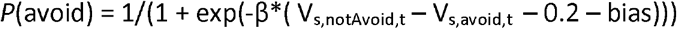

where the free parameter *β* determines how driven *P*(avoid) is by the difference in value between the two possible actions (*V_s,notAVoid,t_* – *V_s,avoid,t_*), and the *bias* parameter determines overall bias towards a particular action (avoiding or not avoiding).

For both samples, models were fit to choice (avoidance) data using the variational Bayes approach to model inversion implemented in the VBA toolbox (Daunizeau et al., 2014; available at mbb-team.github.io/VBA-toolbox), run in Matlab. Model fit was performed in a mixed-effects framework. Simply, after the first round of model inversion, the individual posterior free parameter value estimates are used to approximate the population distribution these values were drawn from, which is then used as prior for the next round of inference, until convergence (no further group-level reduction free energy). This approach reduces the likelihood of outliers in any individual parameter estimates.

Model comparison was by random-effects Bayesian model comparison (Rigoux et al., 2014). This method of model comparison assumes that the population is composed of subjects that differ in terms of the model that describes them best, then induces a hierarchical probabilistic model that can be inverted to derive the posterior density over model frequencies, given participants’ data. Under this approach, the critical metric for any given model is its exceedance probability, or the likelihood that that particular model is more frequent than all other models in the comparison set.

### Functional imaging data Pre-processing

Functional imaging data were pre-processed using SPM12 (Wellcome Trust Centre for Neuroimaging, www.fil.ion.ucl.ac.uk/spm) in Matlab. Briefly, functional images were realigned to the first functional image in each sequence, unwarped, corrected for time of acquisition, and normalized to MNI space via tissue probability maps derived from the co-registered structural image. The full pre-processing pipeline available is available at osf.io/f9drs as a BIDS-compatible Matlab script (Gorgolewski et al., 2016). Finally, images were smoothed via convolution with an 8mm full-width at half-maximum Gaussian kernel for the univariate (but not multivariate) analysis.

Breathing belt data were processed using the PhysIO toolbox (Kasper et al., 2017; available at translationalneuromodeling.org/tapas), which provides physiological noise correction for functional imaging data using the Fourier expansion of respiratory phase implemented in the RETROICOR algorithm (Glover et al., 2000).

### Univariate analysis

Functional imaging data were first analysed according to a mass univariate approach based on the general linear model for time series data in each voxel, as implemented in SPM12. This enables detection of whether variance in BOLD in each voxel is significantly related to modelled internal quantities (i.e. if particular model terms are encoded in BOLD signal time course), with relative spatial specificity. Several models were fit to individual BOLD time series data using restricted maximum likelihood estimation to produce individual statistical maps at the 1^st^ level, which were used to determine significance at the 2^nd^ level using one-sample t-tests in a random-effects framework.

All first level models included the following regressors of no interest: 8 respiration and 6 movement regressors (with translation >1.5mm or rotation >1⍰ on any trial resulting in the inclusion of an additional outlier regressor), plus delta functions at the time of shock receipt. In addition:

*Model 1: Expected value analysis*. For ease of interpretability, modelled internal value of *not* avoiding on a given trial (*V_s,avoid,t_*) was multiplied by -1 to effectively represent predicted *P*(shock) for that particular stimulus. The imaging model consisted of delta functions for CS onset (all trials), with parametric modulators of (i) estimated *P*(shock) according to the perceptual only model (ii) estimated P(shock) according to the perceptual + value-based generalization model.

*Model 2: Prediction error analysis*. Prediction error (PE) was defined as the difference between predicted and actual outcome on a given trial, or (*R_t_ – V_s,a,t_)*. NB by definition this is equal to 0 on all trials where an avoidance response was made. The imaging model consisted of delta functions at the time of expected outcome delivery (all trials), with parametric modulators of (i) trial PE according to the perceptual only model (ii) trial PE according to the perceptual + value-based generalization model. Here, positive PEs represent shock receipt (where predicted P(shock) was <1), and negative PEs represent shock omission (where predicted P(shock) was >0).

All regressors were convolved with a canonical haemodynamic response function, with correction for low-frequency drift using high pass filtering (1/128s) and correction for serially correlated errors by fitting of a first-order autoregressive process (AR(1)).

Computational model-based regressors were derived using individual subject free parameter values, and all regressors were orthogonalised during model estimation. SPM assigns variance to parametric modulators in a successive fashion, such that in an orthogonalised framework, a significant finding from a second parametric modulator represents that due to variance over and above that which has been assigned to the first modulator (Mumford et al., 2015). First level models did not include nuisance regressors at time of avoidance responses, as these were highly collinear with trial-by-trial expected value estimates (by definition, since these are derived from avoidance behaviour). Therefore we would expect some motor preparation responses to be included in the EV(not avoid)/predicted P(shock) contrast in Model 1. This problem should be less evident when examining trial-by-trial PEs (Model 2), as this considers BOLD signal at the time of outcome rather than during the decision period.

An initial cluster-forming threshold of p<0.001 (uncorrected), cluster size ≥10, was applied to 2nd level SPMs, followed by cluster-level family wise error (FWE) rate correction at the whole-brain level (*p*_WB_). Small-volume correction (*p*_SVC_) was applied in *a priori* regions of interest (ROIs): namely the insula, amygdala, striatum, primary visual cortex (V1) and ventromedial prefrontal cortex (vmPFC) (see main text). ROIs were defined anatomically using the automatic anatomical labelling (aal) atlas (Tzourio-Mazoyer et al., 2002) in SPM (‘striatum’ = caudate + putamen + pallidum; ‘V1’=Brodmann Area 17; ‘vmPFC’=medial orbitofrontal cortices).

Only voxels present in all subjects were included in the analysis. For display purposes, statistical maps were thresholded at *p*<0.001 (uncorrected), and overlain on a high quality mean MNI-space structural image available as part of the MRIcroGL package. All quoted voxel coordinates refer to MNI space, in mm.

### Multivariate analysis

Representational similarity analysis (RSA) was carried out using materials from the RSA toolbox (Nili et al., 2014; available at github.com/rsagroup/rsatoolbox), run in Matlab.

For this analysis, pre-processed (but unsmoothed) time series data extracted from all voxels of each ROI were first multivariately noise normalized. We calculated linear discriminant contrast (LDC; Walther et al., 2015) values between pairs of stimulus categories (CS-, GS, CS+) as a robust estimate of representational dissimilarity. This approach involves construction of an optimal decision boundary (hyperplane) between pairs of multivariate representations (i.e. BOLD signal in all voxels, see **Figure 4a**). LDC values are a continuous measure of representational distance (dissimilarity) drawn by sampling of a dimension orthogonal to this decision boundary (Fisher’s linear discriminant). To ensure distances were unbiased by noise (and therefore had a meaningful zero point), LDC values were estimated using a leave-one-out cross-validation approach across functional imaging runs (this constitutes a cross-validated estimate of the Mahlanobis distance, Waltheret al., 2015).

*A priori* regions of interest were the same as for the univariate analysis. However, per our analysis plan, where possible anatomical ROIs were replaced by functional ROIs defined from the group-level univariate analysis. Specifically, the anterior insula and caudate clusters identified in **Figure 3b** were substituted for whole structure anatomical ROIs. This was done on the basis that 1) the univariate analysis indicated involvement of these voxels in specific value-related generalization processes, and 2) previous analysis has shown that reliability of LDC RDMs falls off sharply for larger ROIs (>~250 voxel, Walther et al., 2015; anatomical ROIs for whole insula = 1019 voxels, for whole striatum= 3482 voxels; functional anterior insula ROI= 96 voxels, functional caudate ROI=64 voxels).

### Questionnaire data

Questionnaire total and individual item scores were feature scaled (z-scored across participants) prior to further analysis.

Factor analysis was carried out as described in Gillan et al. (Gillan et al., 2016), implemented in R v3.4.0 (R Foundation for Statistical Computing), using the factanal function (psych package) with oblique (oblimin) rotation. The number of factors to extract was determined using the Cattell-Nelson-Gorsuch (Gorsuch and Nelson, 1981) method (nFactors package), whereby successive scree plot gradients are analysed to determine the “elbow” point after which there is little gain in retaining additional factors. Factor names were chosen on the basis of the highest-loading items for each factor.

### Individual differences

Normality of distribution of individual variables (or within-subject differences in variables) was assessed using the Shapiro-Wilk test, and where appropriate, non-parametric statistics were employed for pairwise tests.

Individual model parameters governing value-based generalization (⍰_A_/⍰_N_) were related to variables of interest (multivariate representational similarity in the fMRI sample, self-reported psychopathology in online sample) using weighted least squares regression. This method produces the maximum likelihood regression estimate when noise is not constant across measurements (i.e. data are heteroscedatic; Carroll and Ruppert, 1988). As the VBA toolbox yields the variance of posterior parameter estimates as well as the mean, weights were defined as the precision of individual parameter estimates (i.e. 1/posterior variance).

Analysis was implemented in R using the function lm (psych package). Age (z-scored) and gender (binary scored as male *vs* female/other) were included in all questionnaire data models as predictors of no interest. In R syntax:

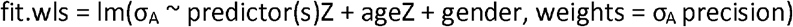

For comparison of brain data to overall GS avoidance in the fMRI sample, we used non-weighted models, as precision estimates were not available. Mean avoidance across different trial types was z-scored within-participants, in order to gain a measure of *relative* GS avoidance (i.e. taking into individual variation in tendency to avoid on CS- and CS+ trials).

Where candidate predictors were significantly collinear (as was the case for the questionnaire data), predictors were implemented in separate regression models (as per Gillan et al). Multiple comparisons correction for these models was achieved via the Nyholt-Bonferroni correction proposed by Li and Ji (2005), which yields a modified Bonferroni correction for non-independent (related) variables by estimating the ‘effective number of independent variables’ from the eigenvalues of their correlation matrix.

As a more robust test, we complemented these linear regression models with crossvalidated regularized regression models, where all predictors were included in a single model. Specifically, we used least absolute shrinkage and selection operator (LASSO) regression (Tibshirani, 1996) with leave-one-out cross-validation. This approach effectively shrinks non-significant predictors to zero, and provides a more robust estimate of regression coefficients. This was implemented using the glmnet package in R.In R syntax:

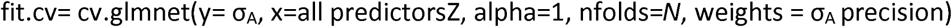

## Supplemental Information

### Supplemental Results

#### Time-on-task analysis of avoidance behaviour

In both groups of participants, there were significant effects of both CS type and block number, and a CS type*block interaction, on proportionate avoidance responding (fMRI: *F*_2,50_ = 406.3, *F*_4,100_=6.14, *F*_8,200_=8.68, respectively; AMT: *F*_2,962_=1077.9, *F*_4,196_2=24.3, *F*_8,3848_=263.0, respectively; all *p*<0.001, repeated-measures ANOVA). In the fMRI sample, the CS type*block interaction was driven by lower avoidance for CS+ stimuli in block 1 compared to the rest of the task (*p*≤0.004; other CS types no significant differences between blocks; pairwise comparisons Bonferroni corrected for multiple comparisons). This suggests a strategy of exploratory non-avoidance to enable proper learning of CS+ stimuli in block 1, but fairly constant generalization of avoidance across later blocks. In the AMT sample, there was also lower avoidance for CS+ stimuli in block 1 vs other blocks (all *p*<0.001), but also decrease in avoidance for CS- stimuli in later blocks (3-5) vs earlier blocks (1 and 2; both *p*<0.001). Overall GS avoidance showed small increases then decreases over first 3 blocks (*p*<0.001), before stabilising between blocks 4 and 5 (p>0.5, Bonferroni-corrected pariwise comparisons; see **Figure S2**).

#### Univariate fMRI analysis with changing perceptual acuity

Univariate fMRI results from a model where participants’ perceptual acuity on the generalization task was modelled as a linear function between their pre- and postconditioning measurements were consistent with results from the model where perceptual discriminability of GSs was held constant at 75%. Specifically, there was signficant positive encoding of additional variance in predicted P(shock) from the value-based model in the anterior insula, bilaterally (left: *p*_WB_=0.005, *k*=87, peak voxel [-27,23,-1], *Z*=5.17; right: *p*_WB_=0.012, *k*=73, peak voxel [33,23,2], *Z*=4.60), mid cingulate cortex (supplementary motor area; *p*_WB_=0.008, *k*=79, peak voxel [9,14,44], *Z*=4.97), and the right caudate nucleus (*p*_SVC_=0.026, *k*=21, peak voxel =[12,8,8], *Z*=3.96). There was also significant negative encoding of additional variance in trial prediciton error signal from the value-based model in the right anterior insula (*p*_SVC_=0.024, *k*=18, peak voxel [48,5,2], *Z*=3.93) and left putamen (*p*_SVC_=0.014, *k*=24, peak voxel [-27,-7,2], *Z*=3.69).

**Figure S1.**
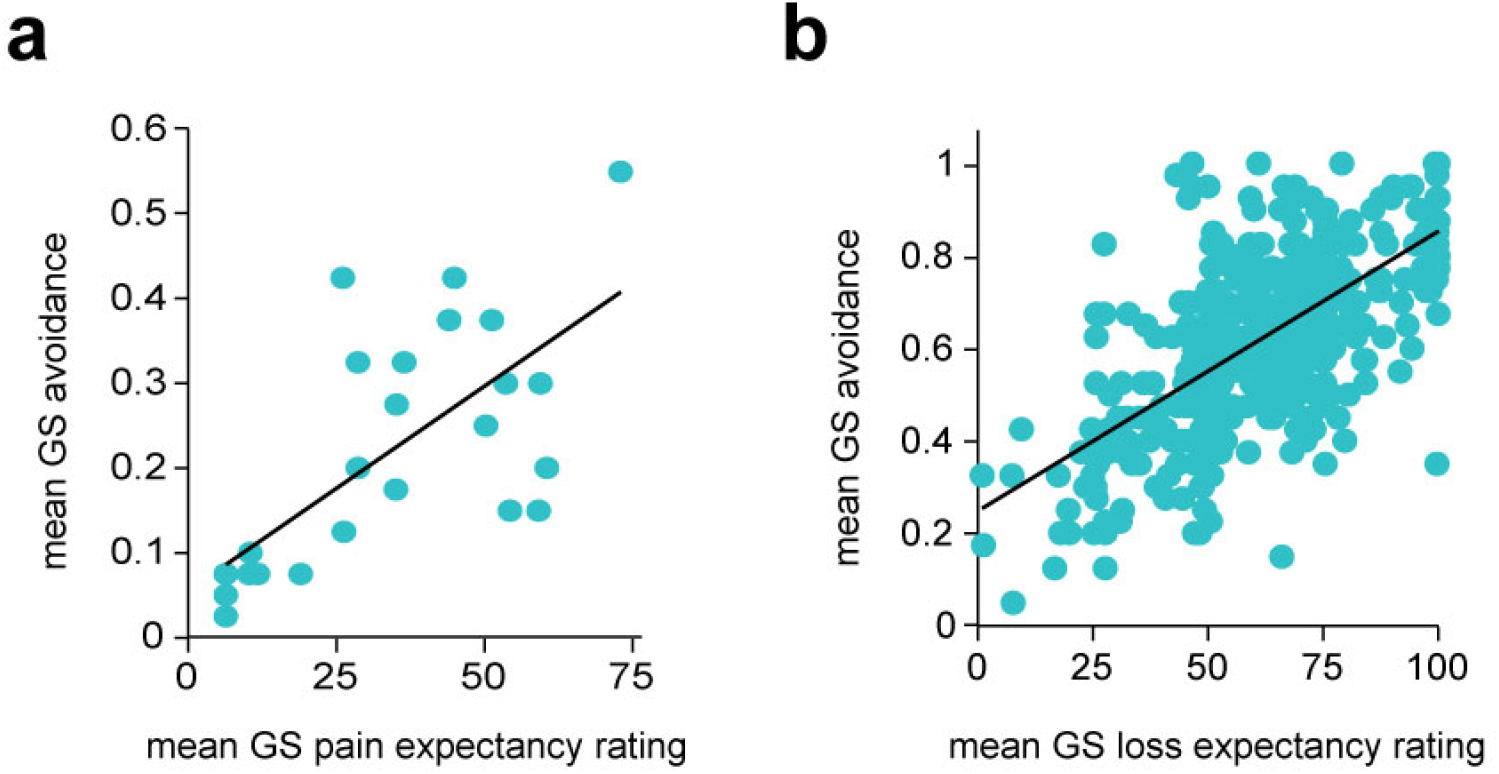
Relationship between mean avoidance on generalization stimulus (GS) trials during the generalization of instrumental avoidance task, and mean post-task visual analogue scale pain/loss expectancy ratings. **a**, fMRI, **b**, AMT, samples (Spearman’s ρ= 0.692, 0.641, respectively).

**Figure S2.**
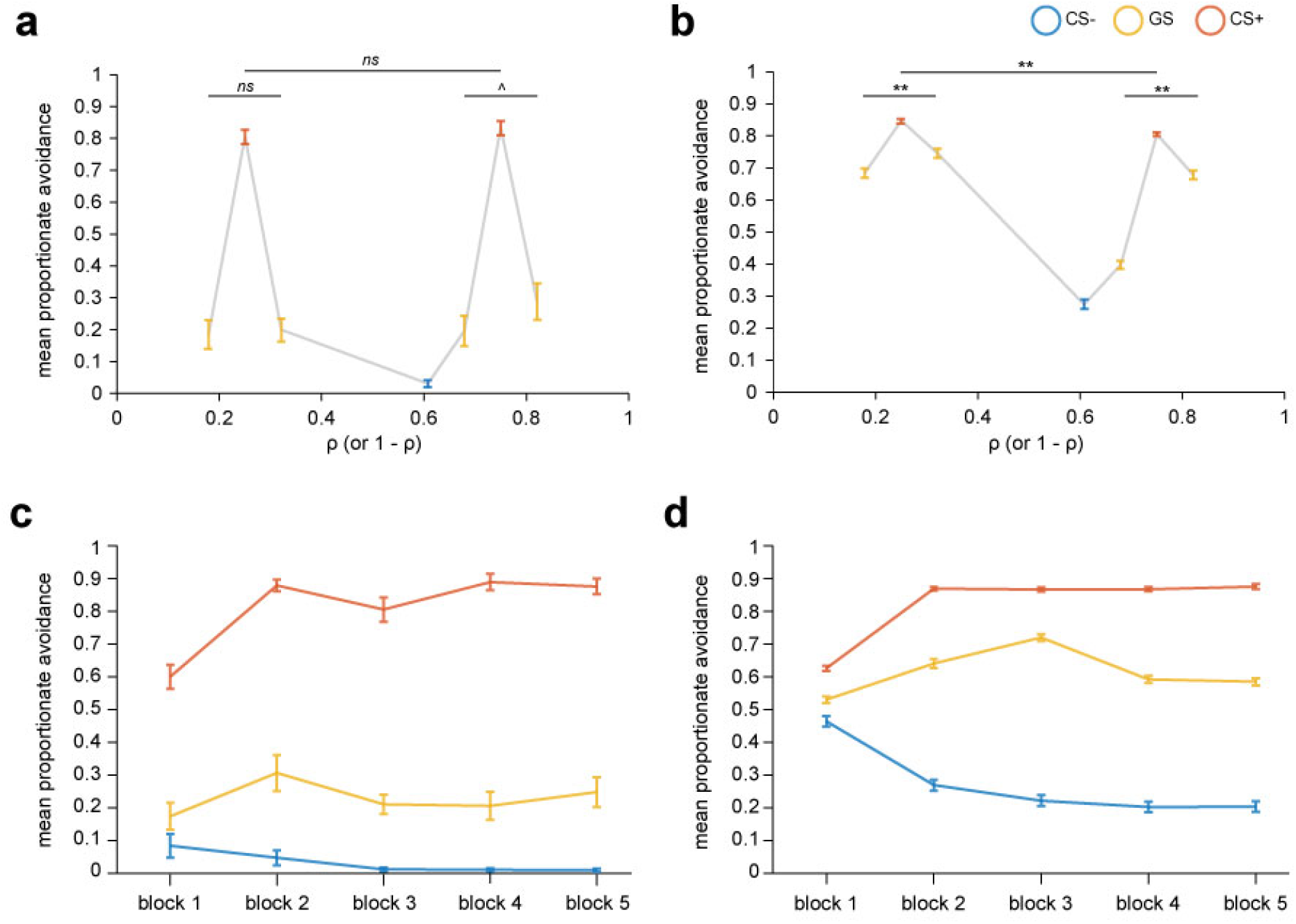
Proportionate avoidance for individiual task stimuli (top row) and by CS type and block number (bottom row) for the generalization of instrumental avoidance task. **a**, fMRI, **b**, AMT, samples, *ns, p*>0.3. ^*p*=0.19, \*\**p*<0.001, repeated-measures ANOVA for differences in mean avoidance across generalization stimuli (GSs). **c**, fRMI, **d**, AMT, samples. Error bars represent standard error.

**Figure S3.**
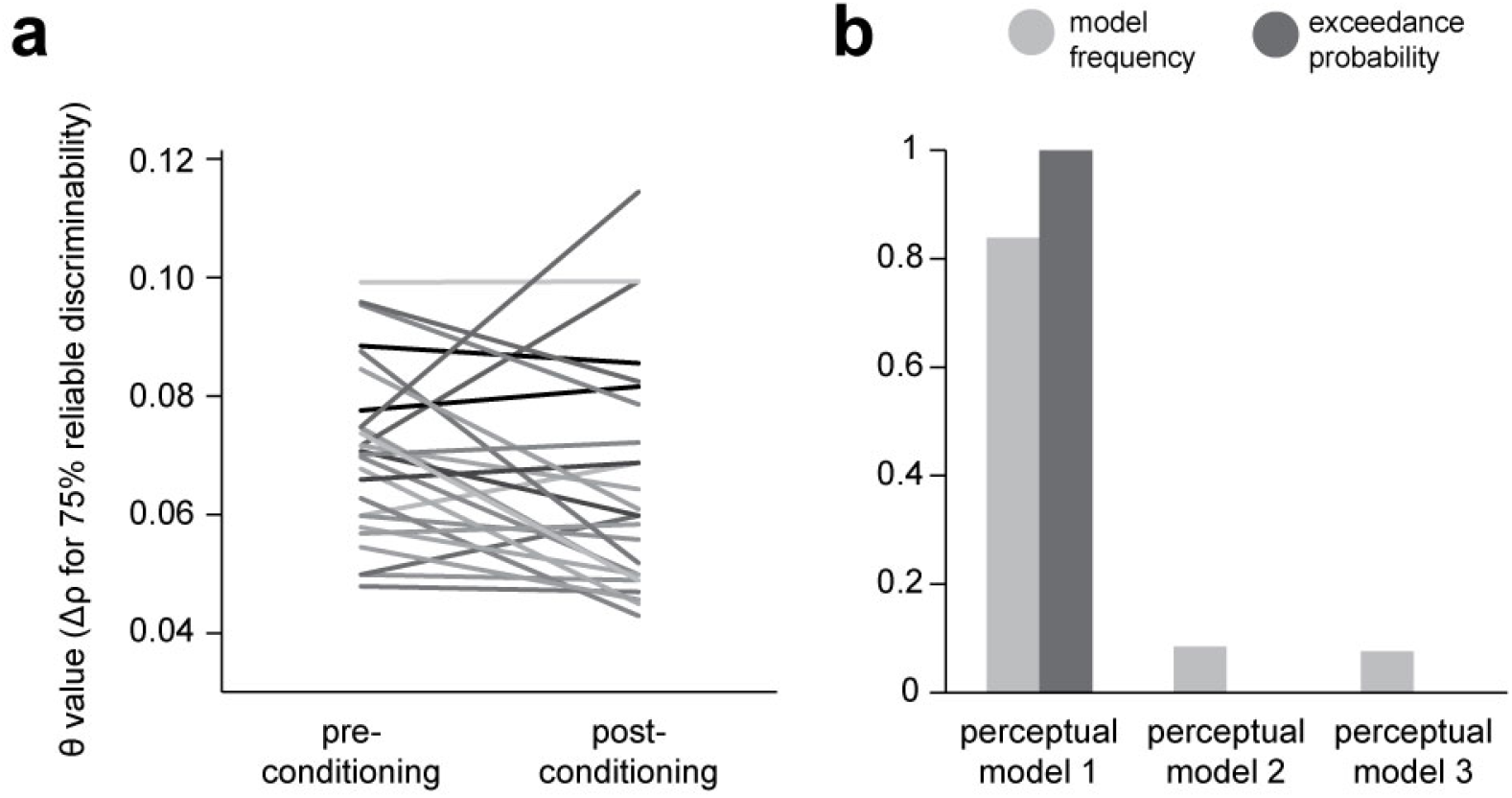
Effects of conditioning on perceptual acuity for task stimuli. **a**, Changes in perceptual acuity as measured by the delayed-punished perceptual task, before and after conditioning (performance of the generalization of instrumental avoidance task), for each participant in the fMRI group. θ, change in stimulus ‘spikiness’ parameter p required to identify a shape as different on 75% of trials. **b,** Results of Bayesian model comparison carried out to determine the best model of participants’ perceptual performance during the generalization of instrumental avoidance task. Model 1, a perceptual-only generalization model in which perceptual discriminability of GSs is fixed at 75%. Model 2, a perceptual-only generalization model in which perceptual discriminability of GSs is fixed at the value determined by the post-conditioning acuity test. Model 3, a perceptual-only generalization model in which GS discriminability changes linearly from the pre to post-conditioning derived value, over the course of the task. Model frequency, proportion of participants for whom a model was the best model; exceedance probability, probability that the model in question is the most frequently utilized in the population

**Figure S4.**
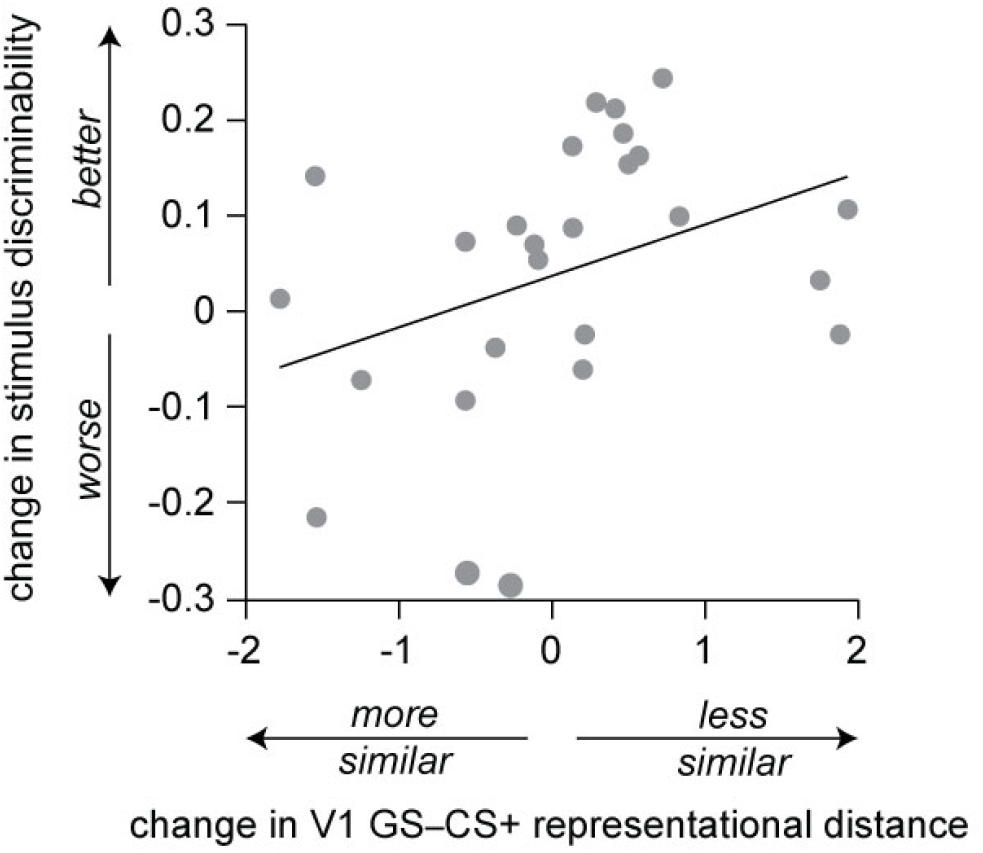
Relationship between change in stimulus discriminability, pre vs postconditioning, and change in GS–CS+ representational distance (CV LDC) in the primary visual cortex (V1) over the course of the generalization task. Pre-conditioning (day 1 testing), discriminability for target stimulus ± θ was 0.75 (75% correct difference judgments), by definition. Post-conditioning (day 3 testing), mean discriminability for target ± θ was 0.79 (SD 0.14).

**Table S1.** Demographic information for study participants. Unless otherwise specified, figures represent mean (SD). STAI, Spielberger State-Trait Anxiety Inventory (trait score only); AMI, Apathy Motivation Index; OCI-R, Obsessive-Compulsive Index (Revised); PHQ9, Physician’s Health Questionnaire 9 (a brief measure of mood disorder symptoms); BIS-11, Barratt Impulsivity Scale (version 11); CSQ global, Cognitive Style Questionnaire (short-form) ‘cognitive globalisation’ subscale.

|  | fMRI sample<br>(N=26) | AMT sample<br>(N=482) | Possible<br>range |
| --- | --- | --- | --- |
| Age | 25.3 (5.6) | 37.2 (11.4) | - |
| Gender (%F) | 13 (50%) | 249 (52%) | - |
| STAI total | 42.0 (8.1) | 39.9 (12.8) | 20-80 |
| AMI total | - | 40.1 (8.5) | 0-72 |
| OCI-R total | - | 11.3 (10.8) | 0-72 |
| PHQ9 total | - | 4.3 (5.1) | 0-24 |
| BIS-11 total | - | 56.6 (10.8) | 30-120 |
| CSQ global | - | 29.1 (7.9) | 0-48 |

**Table S2.**
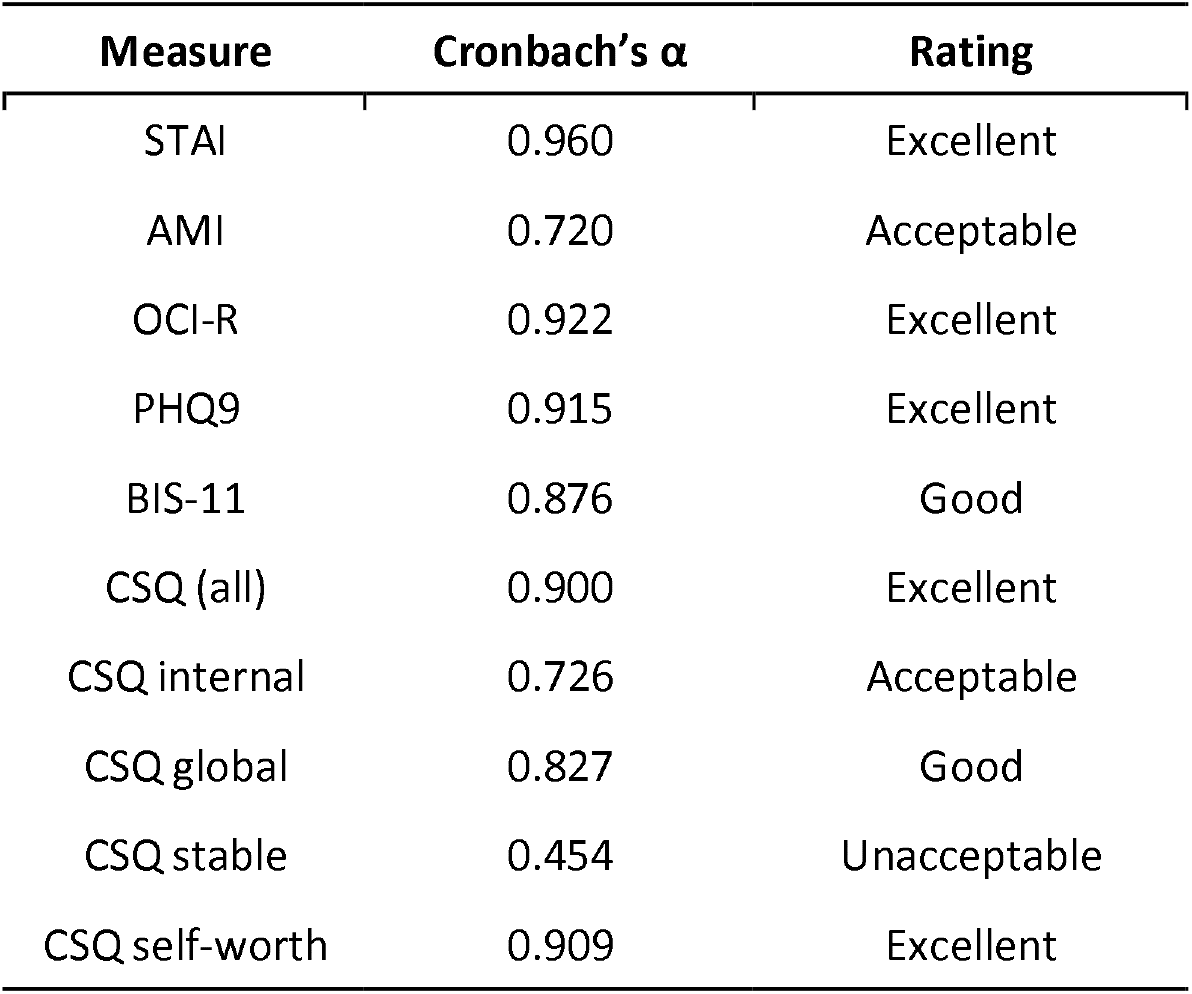
Internal reliability of questionnaire scores in the AMT sample. STAI, Spielberger State-Trait Anxiety Inventory (trait score only); AMI, Apathy Motivation Index; OCI-R, Obsessive-Compulsive Index (Revised); PHQ9, Physician’s Health Questionnaire 9 (a brief measure of mood disorder symptoms); BIS-11, Barratt Impulsivity Scale (version 11); CSQ, Cognitive Style Questionnaire (short-form).

**Table S3a.** Individual item loadings onto Factor 1 – “Intrusive thoughts and anxiety”. (above threshold of ± 0.25). Text in square brackets is to aid interpretation of reverse-scored items.

| Item | Loading |
| --- | --- |
| I am upset by unpleasant thoughts that come into my mind against my will | 0.762 |
| I find it difficult to control my thoughts | 0.749 |
| I frequently get nasty thoughts and have difficulty in getting rid of them | 0.725 |
| I get upset if others change the way I have arranged things | 0.682 |
| I check things more often than necessary | 0.673 |
| I get upset if things are not arranged properly | 0.632 |
| I have disturbing thoughts | 0.627 |
| I need things to be arranged in a particular order | 0.624 |
| I get in a state of tension or turmoil as I think over my recent concerns and interests | 0.619 |
| I have racing thoughts | 0.605 |
| Little interest or pleasure in doing things | 0.598 |
| I repeatedly check doors, windows, drawers etc. | 0.593 |
| I sometimes shave or wash or clean myself simply because I feel contaminated | 0.591 |
| Some unimportant thought runs through my mind and bothers me | 0.588 |
| I worry too much over something that doesn't really matter | 0.575 |
| I feel that difficulties are piling up so that I cannot overcome them | 0.574 |
| Feeling down, depressed, or hopeless | 0.569 |
| I feel I have to repeat certain numbers | 0.564 |
| I have saved up so many things that they get in the way | 0.562 |
| I avoid throwing things away because I am afraid I might need them later | 0.562 |
| I wash my hands more often and longer than necessary | 0.558 |
| I often have extraneous or intrusive thoughts when trying to think | 0.557 |
| Feeling tired or have little energy | 0.555 |
| I wish I could be as happy as others seem to be | 0.550 |
| Trouble concentrating on things such as reading the newspaper or watching television | 0.550 |
| I collect things I don't need | 0.545 |
| Poor appetite or over-eating | 0.541 |
| I feel nervous and restless | 0.540 |
| I feel compelled to count while I am doing things | 0.540 |
| Feeling bad about yourself or that you are a failure or have let yourself down | 0.539 |
| I take disappointments so keenly I can't put them out of my mind | 0.534 |
| I find it difficult to touch an object I know has been touched by a stranger | 0.507 |
| I feel like a failure | 0.491 |
| I feel like there are good and bad numbers | 0.490 |
| I repeatedly check gas/water taps and light switches after turning them off | 0.470 |
| I feel inadequate | 0.466 |
| Trouble falling or staying asleep, or sleeping too much | 0.465 |
| Moving or speaking so slowly that other people could have noticed (or the opposite) | 0.429 |
| I am [un]happy | 0.381 |
| I am [not] content | 0.381 |
| I lack self-confidence | 0.370 |
| I [do not] feel secure | 0.362 |
| I feel [un]pleasant | 0.345 |
| I feel [un]satisfied with myself | 0.337 |
| I feel [un]rested | 0.332 |
| I am [not] 'cool, calm, and collected' | 0.316 |
| I squirm during plays or lectures | 0.307 |
| I am a[n un]steady person | 0.296 |
| I am restless at the theatre or in lectures | 0.286 |
| I [don't] like to think about complex problems | -0.264 |
| Getting on badly with my parents is [not] the fault of other people or circumstances | -0.275 |
| I am happy-go-lucky | -0.294 |

**Table S3b.** Individual item loadings onto Factor 2 – “Low self-worth”. (above threshold of ± 0.25). Text in square brackets is to aid interpretation of reverse-scored items.

| Item | Loading |
| --- | --- |
| That people were not interested in me at the party [says something] about me as a person | 0.742 |
| Getting a negative reaction to my talk [says something] about me as a person | 0.732 |
| People not being interested in me at the party means there is something wrong with me as a person | 0.700 |
| The reason that the talk went badly [will] stop me from enjoying other things | 0.694 |
| I will [not] start afresh in the future and forget about the reason people were not interested in me | 0.659 |
| That I'm not in a romantic relationship now [says something] about me as a person | 0.651 |
| The reason that this evaluation went badly [will] stop be from enjoying other things | 0.642 |
| Getting a negative evaluation on my job performance [says something] about me as a person | 0.632 |
| I will [not] start afresh in the future and forget about the reason for the talk going badly | 0.625 |
| Doing badly on this talk means that there is something wrong with me as a person | 0.593 |
| That I'm not able to get all my work done at the moment [says something] about me as a person | 0.582 |
| I will [not] start afresh in the future and forget about the reasons for my evaluation being negative | 0.572 |
| I will [not] start afresh in the future and forget about the reason I think I'm not in a relationship at the moment | 0.570 |
| The reason why I think other people did not seem interested in me [will] stop me from enjoying other things | 0.558 |
| Doing badly on this evaluation means that there is something wrong with me as a person | 0.544 |
| The reason why I think I can't get my work done [will] stop me from enjoying other things | 0.539 |
| Getting along badly with my parents [says something] about me as a person | 0.532 |
| That I am not in a romantic relationship right now means there is something wrong with me as a person | 0.531 |
| It is my fault that people are not interested in me | 0.526 |
| The reason I am getting on badly with my parents [will] stop me from enjoying other things | 0.503 |
| I will [not] try to start afresh in the future and forget about the reason I think I can't complete my work at the moment | 0.488 |
| That I am not able to complete the work means that there is something wrong with me as a person | 0.486 |
| The reason no one was interested in me causes me problems in all areas of my life | 0.475 |
| The reason I did badly on my talk causes problems for me in all areas of my life | 0.458 |
| I lack self-confidence | 0.447 |
| The reason why I think I'm not in a relationship [will] stop me from enjoying other things | 0.413 |
| Getting along badly with my parents means there is something wrong with me as a person | 0.407 |
| It is my fault the class reacted negatively to my talk | 0.399 |
| People not being interested in me is [not] the fault of other people | 0.386 |
| I feel inadequate | 0.381 |
| The reason causing me not to be in a relationship causes me problems in all areas of my life | 0.380 |
| The reason I did badly on my evaluation causes me problems in all areas of my life | 0.367 |
| I feel [un]satisfied with myself | 0.347 |
| My parents and I will [not] start afresh in the future and forget about the reason for getting along badly | 0.345 |
| I feel [un]pleasant | 0.331 |
| I am [un]happy | 0.331 |
| I feel like a failure | 0.310 |
| I wish I could be as happy as others seem to be | 0.287 |
| I [do not] feel secure | 0.286 |
| I [don't] enjoy choosing what to do from a range of activities | 0.285 |
| I take disappointments so keenly I can't put them out of my mind | 0.280 |
| I am [not] content | 0.275 |
| I am [not] 'cool, calm and collected' | 0.270 |
| The same reason causing me to not complete my work causes me problems in all areas of my life | 0.262 |
| I [don't] like to think about complex problems | 0.259 |
| It is my fault that I am not in a romantic relationship | 0.258 |
| The negative reaction to my talk is [not] the fault of other people or circumstances | 0.252 |
| I act on the spur of the moment | -0.260 |
| I am happy-go-lucky | -0.325 |
| The reason that I'm not in a relationship now will not prevent me from being in a relationship in the future | -0.459 |
| The reason that I can't get my work done now will not prevent me from being able to get everything done in the future | -0.506 |
| The reason that people didn't seem interested in me at this party will not prevent me from enjoying similar situations in the future | -0.715 |

**Table S3c.** Individual item loadings onto Factor 3 – “Low self-control”. (above threshold of ± 0.25). Text in square brackets is to aid interpretation of reverse-scored items.

| Item | Loading |
| --- | --- |
| I [don't] plan tasks carefully | 0.775 |
| I am [not] a careful thinker | 0.681 |
| I am [not] self-controlled | 0.659 |
| I am [not] a careful thinker | 0.633 |
| I [don't] plan trips well ahead of time | 0.627 |
| I [do not] plan for job security | 0.615 |
| I [do not] concentrate easily | 0.611 |
| I do things without thinking | 0.552 |
| I [do not] save regularly | 0.546 |
| I don't pay attention | 0.512 |
| I act on impulse | 0.506 |
| When I decide to do something I am [not] motivated to see it through to the end | 0.503 |
| I am an [un]steady person | 0.484 |
| I am [not] future oriented | 0.479 |
| I spend more than I earn | 0.447 |
| I act on the spur of the moment | 0.438 |
| I [don't] get things done when they need to be but require [reminders] from others | 0.433 |
| I make decisions [with difficulty] | 0.400 |
| I buy things on impulse | 0.397 |
| I say things without thinking | 0.363 |
| I am [not] 'cool, calm and collected' | 0.360 |
| I get easily bored when solving thought problems | 0.350 |
| I am [not] content | 0.339 |
| When I have something I need to do I [don't] do it straight away so it is out of the way | 0.334 |
| When I decide to do something I am [not] able to make an effort easily | 0.327 |
| I feel [un]satisfied with myself | 0.323 |
| I often have extraneous or intrusive thoughts when trying to think | 0.318 |
| I feel [un]pleasant | 0.312 |
| I [do not] feel secure | 0.305 |
| I feel [un]rested | 0.298 |
| If I realise I have been unpleasant to someone I will [not] feel terribly guilty afterwards | 0.276 |
| I am [un]happy | 0.273 |
| I change jobs | 0.269 |
| Trouble concentrating on things such as reading the newspaper or watching television | 0.267 |
| I [do not] like puzzles | 0.265 |
| I [don't] enjoy choosing what to do from a range of activities | 0.264 |
| I [don't] like to think about complex problems | 0.261 |
| I get upset if others change the way I have arranged things | -0.292 |
| I need things to be arranged in a particular order | -0.297 |
| I get upset if things are not arranged properly | -0.310 |

